# TDM: a temporal decomposition method for removing venous effects from task-based fMRI

**DOI:** 10.1101/868455

**Authors:** Kendrick Kay, Keith W. Jamison, Ruyuan Zhang, Kamil Ugurbil

**Affiliations:** Center for Magnetic Resonance Research (CMRR), Department of Radiology, University of Minnesota

**Keywords:** high-resolution fMRI, veins, vasculature, hemodynamic timecourse, hemodynamic response function, primary visual cortex, eccentricity, 7T

## Abstract

Most functional magnetic resonance imaging (fMRI) is conducted with gradient-echo pulse sequences. Although this yields high sensitivity to blood oxygenation level dependent (BOLD) signals, gradient-echo acquisitions are heavily influenced by venous effects which limit the ultimate spatial resolution and spatial accuracy of fMRI. While alternative acquisition methods such as spin-echo can be used to mitigate venous effects, these methods lead to serious reductions in signal-to-noise ratio and spatial coverage, and are difficult to implement without leakage of undesirable non-spin-echo effects into the data. Moreover, analysis heuristics such as masking veins or sampling inner cortical depths using high-resolution fMRI may be helpful, but sacrifice information from many parts of the brain. Here, we describe a new analysis method that is compatible with conventional gradient-echo acquisition and provides venous-free response estimates throughout the entire imaged volume. The method involves fitting a low-dimensional manifold characterizing variation in response timecourses observed in a given dataset, and then using identified early and late timecourses as basis functions for decomposing responses into components related to the microvasculature (capillaries and small venules) and the macrovasculature (veins), respectively. We show that this *Temporal Decomposition through Manifold Fitting (TDM)* method is robust, consistently deriving meaningful timecourses in individual fMRI scan sessions. Moreover, we show that by removing late components, TDM substantially reduces the superficial cortical depth bias present in gradient-echo BOLD responses and eliminates artifacts in cortical activity maps. TDM is general: it can be applied to any task-based fMRI experiment, can be used with standard- or high-resolution fMRI acquisitions, and can even be used to remove residual venous effects from specialized acquisition methods like spin-echo. We suggest that TDM is a powerful method that improves the spatial accuracy of fMRI and provides insight into the origins of the BOLD signal.

## 1. Introduction

Among the handful of noninvasive techniques that permit the study of human brain activity, functional magnetic resonance imaging (fMRI) based on blood oxygenation level dependent (BOLD) contrast has emerged as by far the most commonly used approach and has proven to be an extremely useful tool in cognitive neuroscience. A primary advantage of fMRI over other measurement techniques that report on neural activity (e.g., EEG, MEG, PET) is its spatial resolution. Whole-brain volumes sensitized to the BOLD effect are routinely acquired at a resolution of 2–3 mm, and even at sub-second imaging times as exemplified by data from the Human Connectome Project (Ugurbil et al., 2013). However, efforts to increase the spatial resolution of fMRI further—especially to reach the sub-millimeter scale of mesoscopic brain organization—face a major challenge imposed by the indirect nature of fMRI signals. The BOLD response is mediated by neurovascular coupling (Attwell and Iadecola, 2002) that links neuronal activity to secondary hemodynamic and metabolic alterations. An undesirable consequence of this mechanism is the “draining vein” confound that affects the most commonly employed gradient-echo BOLD fMRI technique, first noted early in the history of fMRI (Menon et al., 1993) and investigated in numerous subsequent studies. Deoxyhemoglobin changes originating at the site of neuronal activity subsequently propagate through venules and veins; these effects may appear as activation displaced from the original site of neural activity by as much as 4 mm (Turner, 2002) and may reflect neural activity pooled over large spatial scales, thus degrading spatial specificity with respect to underlying neural activity patterns (Bianciardi et al., 2011; Kay et al., 2019; Olman et al., 2007; Shmuel et al., 2007). In response to this problem, the field has long sought to measure BOLD responses from the microvasculature (e.g. capillaries and venules) while avoiding BOLD responses from the macrovasculature (e.g. veins) (Cheng, 2018; Ugurbil, 2016; Yacoub and Wald, 2018). The problem of the macrovasculature is especially critical to resolve given the growing popularity in the neuroscience community of using fMRI to probe laminar-specific responses (see De Martino et al., 2018; Dumoulin et al., 2018; Lawrence et al., 2017 for reviews).

In order to avoid the specificity loss caused by veins, the field has traditionally turned to the use of spin-echo acquisition (e.g. Parkes et al., 2005; Yacoub et al., 2008) instead of conventional gradient-echo acquisition. However, spin-echo involves increased energy deposition, longer volume acquisition times, and lower signal-to-noise ratio. Thus, in order to maintain measurement sensitivity, the experimenter is generally forced to reduce spatial coverage and/or substantially increase the amount of data collected per experimental condition. These unappealing options are often dealbreakers for neuroscientists, given that measuring multiple brain regions is often critical, the sensitivity of fMRI is already relatively low to start with, and increasing the duration of an experiment beyond more than a factor of two or so is often impractical.

Here, we introduce an analysis method, called *Temporal Decomposition through Manifold Fitting (TDM)*, that identifies and removes venous-related signals from task-based fMRI data. The TDM method is simple, principled, and is compatible with a variety of experimental protocols including those based on gradient-echo acquisitions. We demonstrate TDM on visual experiments conducted on human subjects, and show that TDM consistently removes unwanted venous effects while maintaining a reasonable level of sensitivity. We end with a discussion of the strengths and limitations of the TDM method and a comparison of TDM to other approaches for reducing venous effects in fMRI. Code implementing TDM is freely available at https://osf.io/j2wsc/.

## 2. Materials and Methods

### 2.1. Subjects

Eleven subjects (five males, six females; one subject, S1, was an author (K.K.)) participated in the experiments described in this study. All subjects had normal or corrected-to-normal visual acuity. Informed written consent was obtained from all subjects, and the experimental protocol was approved by the University of Minnesota Institutional Review Board.

We conducted four experiments. Experiment E1 measured responses to eccentricity stimuli using a high-resolution (7T, 0.8 mm) gradient-echo protocol. Experiment E2 measured responses to category stimuli also using the high-resolution gradient-echo protocol; data from this experiment are the same as described in a previous publication (Kay et al., 2019). Experiment E3 measured responses to the same eccentricity stimuli in E1 but used a spin-echo protocol (7T, 1.05 mm). Experiment E4 measured responses to the same eccentricity stimuli in E1 but used a low-resolution (3T, 2.4 mm) gradient-echo protocol.

A total of sixteen datasets (scan sessions) were collected: five corresponding to Experiment E1; seven corresponding to Experiment E2; two corresponding to Experiment E3; and two corresponding to Experiment E4. To facilitate direct comparison, Experiments E3 and E4 were conducted in subjects who also participated in Experiment E1. A full breakdown of subjects, experiments, and datasets is provided in **Figure 1**.

**Figure 1.**
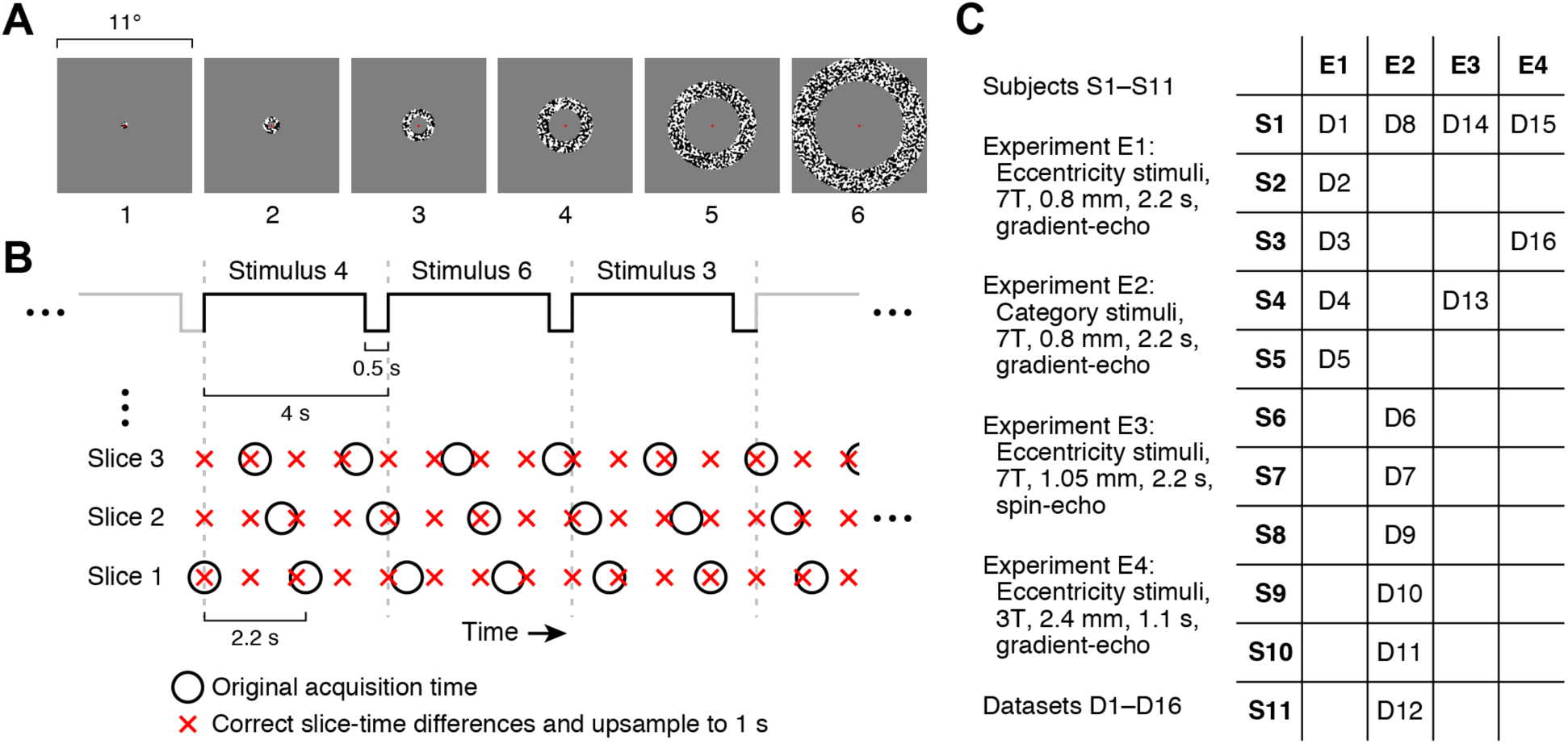
Schematic of experiment. *A*, Eccentricity stimuli. Stimuli consist of six rings varying in eccentricity. A small central dot serves as a fixation point. *B*, Timing of trials and fMRI acquisition. Each stimulus lasts 3.5 s and is followed by a 0.5-s gap before the next trial. Slices are acquired at different times within each 2.2-s TR (black circles). In pre-processing, cubic interpolation is used to resample each voxel’s time-series data to a rate of 1 s such that the same time points are obtained for all voxels (red crosses). Because the trial duration (4 s) is not evenly divisible by the TR (2.2 s), the experiment automatically incorporates jitter between trial onsets and slice acquisition times. *C*, Summary of datasets. Eleven subjects participated in four experiments. Each dataset corresponds to one scan session, and sixteen datasets were collected. (For details on the category stimuli used in Experiment E2, see Section 2.3.)

### 2.2. Stimulus presentation

For the 7T datasets, stimuli were presented using a Cambridge Research Systems BOLDscreen 32 LCD monitor positioned at the head of the scanner bed (resolution 1920 × 1080 at 120 Hz; viewing distance 189.5 cm). For the 3T datasets, stimuli were presented using a NEC NP4100 DLP projector that was focused onto a backprojection screen positioned at the head of the scanner bore (resolution 1024 × 768 at 60 Hz; viewing distance 102 cm). Subjects viewed the monitor or backprojection screen via a mirror mounted on the RF coil. A Mac Pro (7T) or iMac (3T) computer controlled stimulus presentation using code based on Psychophysics Toolbox (Brainard, 1997; Pelli, 1997). Behavioral responses were recorded using a button box.

### 2.3. Experimental design

In the eccentricity experiment (Experiments E1, E3, E4), stimuli consisted of rings positioned at six different eccentricities (i.e. distances from the center of gaze), and were confined to a circular region with diameter 11°. Each ring was filled with a black-and-white contrast pattern that updated at 10 Hz. Ring size scaled with eccentricity, and rings were presented on a neutral gray background (**Figure 1A**). Stimuli were presented in 4-s trials. In a trial, one of the six rings was presented for 3.5 s (35 images presented sequentially, each with duration 0.1 s) and was followed by a brief gap of 0.5 s. Each run lasted 368.116 s and included 12 presentations of each of the 6 rings as well as blank trials (also of 4-s duration). Throughout stimulus presentation, a small semi-transparent dot (50% opacity) was present at the center of the stimulus. The color of the dot switched between red, white, and black every 1–5 s, and subjects were instructed to maintain fixation on the dot and to press a button whenever the color changed. A total of 9 runs were collected in each 7T scan session, and a total of 6 runs were collected in each 3T scan session. (The eccentricity experiment was time-locked to the refresh rate of the LCD monitor, which caused the additional 116 ms in the total run duration. To compensate for this slight offset, we pre-processed the fMRI data for the eccentricity experiment at a sampling rate of 1.000316 s, and then, for simplicity, treated the data in subsequent analyses as if the sampling rate was exactly 1.0 s.)

The category experiment (Experiment E2) was the same as the ‘functional localizer’ experiment conducted in a previous paper (Kay et al., 2019). This experiment (http://vpnl.stanford.edu/fLoc/) was developed by the Grill-Spector lab (Stigliani et al., 2015). Stimuli consisted of grayscale images of different semantically meaningful categories. There were 10 categories, grouped into 5 stimulus domains: characters (word, number), body parts (body, limb), faces (adult, child), places (corridor, house), and objects (car, instrument). Each stimulus was presented on a scrambled background and occupied a square region with dimensions 10° × 10°. Stimuli were presented in 4-s trials. In a trial, 8 images from a given category were presented sequentially, each with duration 0.5 s. Each run lasted 312.0 s and included 6 presentations of each of the 10 categories as well as blank trials (also of 4-s duration). Throughout stimulus presentation, a small red fixation dot was present at the center of the stimulus. Subjects were instructed to maintain fixation on the dot and to press a button whenever they noticed an image in which only the background was present (“oddball” task). A total of 10–12 runs were collected in each scan session.

### 2.4. MRI data acquisition and pre-processing

Acquisition and pre-processing procedures are the same as described in a previous paper (Kay et al., 2019), except for the addition of a spin-echo acquisition protocol. A summary of all procedures is provided below, and we refer the reader to the previous paper for details.

#### 2.4.1. Acquisition

MRI data were collected at the Center for Magnetic Resonance Research at the University of Minnesota. Some data were collected using a 7T Siemens Magnetom scanner equipped with SC72 body gradients and a custom 4-channel-transmit, 32-channel-receive RF head coil. Other data were collected using a 3T Siemens Prisma scanner and a standard Siemens 32-channel RF head coil. Head motion was mitigated using standard foam padding.

Anatomical data were collected at 3T at 0.8-mm isotropic resolution. We used a whole-brain T1-weighted MPRAGE sequence (TR 2400 ms, TE 2.22 ms, TI 1000 ms, flip angle 8°, bandwidth 220 Hz/pixel, no partial Fourier, in-plane acceleration factor (iPAT) 2, TA 6.6 min/scan) and a whole-brain T2-weighted SPACE sequence (TR 3200 ms, TE 563 ms, bandwidth 744 Hz/pixel, no partial Fourier, in-plane acceleration factor (iPAT) 2, TA 6.0 min/scan). Several T1 and T2 scans were acquired for each subject in order to increase signal-to-noise ratio.

Functional data for Experiments E1 and E2 were collected at 7T using gradient-echo EPI at 0.8-mm isotropic resolution with partial-brain coverage (84 oblique slices covering occipitotemporal cortex, slice thickness 0.8 mm, slice gap 0 mm, field-of-view 160 mm (FE) × 129.6 mm (PE), phase-encode direction inferior-superior (F ≫ H in Siemens’ notation), matrix size 200 × 162, TR 2.2 s, TE 22.4 ms, flip angle 80°, echo spacing 1 ms, bandwidth 1136 Hz/pixel, partial Fourier 6/8, in-plane acceleration factor (iPAT) 3, multiband slice acceleration factor 2). Gradient-echo fieldmaps were also acquired for post-hoc correction of EPI spatial distortion (same slice slab as the EPI data, resolution 2 mm × 2 mm × 2.4 mm, TR 391 ms, TE1 4.59 ms, TE2 5.61 ms, flip angle 40°, bandwidth 260 Hz/pixel, no partial Fourier, TA 1.3 min). Fieldmaps were periodically acquired over the course of each scan session to track changes in the magnetic field.

Functional data for Experiment E3 were collected at 7T using spin-echo EPI at 1.05-mm isotropic resolution with partial-brain coverage (64 (or 48 for Dataset D14) slices, slice thickness 1.05 mm, slice gap 0 mm, field-of-view 128 mm (FE) × 111.2 mm (PE), phase-encode direction inferior-superior (F ≫ H in Siemens’ notation; Dataset D13 was reversed H ≪ F), matrix size 122 × 106, TR 2.2 s, TE 39 ms, flip angle 90°, echo spacing 1 ms, bandwidth 1138 Hz/pixel, partial Fourier 6/8, in-plane acceleration factor (iPAT) 2, multiband slice acceleration factor 2). Corresponding gradient-echo fieldmaps were also acquired.

Functional data for Experiment E4 were collected at 3T using gradient-echo EPI at 2.4-mm isotropic resolution with partial-brain coverage (30 slices, slice thickness 2.4 mm, slice gap 0 mm, field-of-view 192 mm (FE) × 192 mm (PE), phase-encode direction anterior-posterior (A ≫ P in Siemens’ notation), matrix size 80 × 80, TR 1.1 s, TE 30 ms, flip angle 62°, echo spacing 0.55 ms, bandwidth 2232 Hz/pixel, no partial Fourier, no in-plane acceleration, multiband slice acceleration factor 2). Corresponding gradient-echo fieldmaps were also acquired.

#### 2.4.2. Pre-processing

T1- and T2-weighted anatomical volumes were corrected for gradient nonlinearities, co-registered, and averaged (within modality). The averaged T1 volume (0.8-mm resolution) was processed using FreeSurfer (Fischl, 2012) version 6 beta (build-stamp 20161007) with the *-hires* option. We generated 6 cortical surfaces spaced equally between 10% and 90% of the distance between the pial surface and the boundary between gray and white matter, increased the density of surface vertices by bisecting each edge, and truncated the surfaces retaining only posterior cortex in order to reduce memory requirements. The resulting surfaces are termed ‘Depth 1’ through ‘Depth 6’ where 1 corresponds to the outermost surface and 6 corresponds to the innermost surface. Cortical surface visualizations were generated using nearest-neighbor interpolation of surface vertices onto image pixels.

Functional data were pre-processed by performing one temporal resampling and one spatial resampling. The temporal resampling consisted of one cubic interpolation of each voxel’s time-series data; this interpolation corrected differences in slice acquisition times and also upsampled the data to 1.0 s (**Figure 1B**). Data were prepared such that the first time-series data point coincides with the acquisition time of the first slice acquired in the first EPI volume. The motivation for upsampling is to exploit the intrinsic jitter between the data acquisition and the experimental paradigm (**Figure 1B**). The spatial resampling consisted of one cubic interpolation of each volume; this interpolation corrected head motion (rigid-body transformation) and EPI distortion (determined by regularizing the fieldmaps and interpolating them over time) and also mapped the functional volumes onto the cortical surface representations (affine transformation between the EPI data and the averaged T2 volume). After pre-processing, the data consisted of EPI time series sampled every 1.0 s at the vertices of the depth-dependent cortical surfaces (Depth 1–6). Finally, for the purposes of identifying vertices affected by venous susceptibility effects, we computed the mean of the EPI time-series data obtained for each vertex and divided the EPI intensities by a fitted 3D polynomial (up to degree 4); this produced bias-corrected EPI intensities that can be interpreted as percentages (e.g. 0.8 means 80% of the brightness of typical EPI intensities).

### 2.5. Data analysis

Code used for data analysis is available at http://github.com/kendrickkay/. The core functions that implement the TDM method are referenced by name in the text below. Sample data and scripts demonstrating the TDM method are available at https://osf.io/j2wsc/.

### 2.6. GLM analysis

We analyzed the pre-processed time-series data using three different GLM models (FIR, Standard, TDM).

The first GLM model, termed FIR (finite impulse response), is a GLM in which separate regressors are used to model each time point in the response to each experimental condition (Dale, 1999). Results from this model are used as inputs to the TDM method. The FIR model characterized the response from 0 s to 30 s after condition onset, yielding a total of 31 regressors for each condition. (Modeling the response to 30 s was sufficient to capture the majority of the hemodynamic responses; see **Figure 5** and **Supplementary Figure 1**.) We divided the trials for each experimental condition into 2 groups using a “condition-split” strategy (Kay et al., 2019), thereby producing two estimates for each response timecourse. Fitting the FIR model produced BOLD response timecourses (timecourses of ‘betas’) with dimensionality *N* vertices × 6 depths × *M* conditions × 31 time points × 2 condition-splits where *N* is the number of surface vertices for a given subject and *M* is the number of conditions in the experiment.

**Figure 2.**
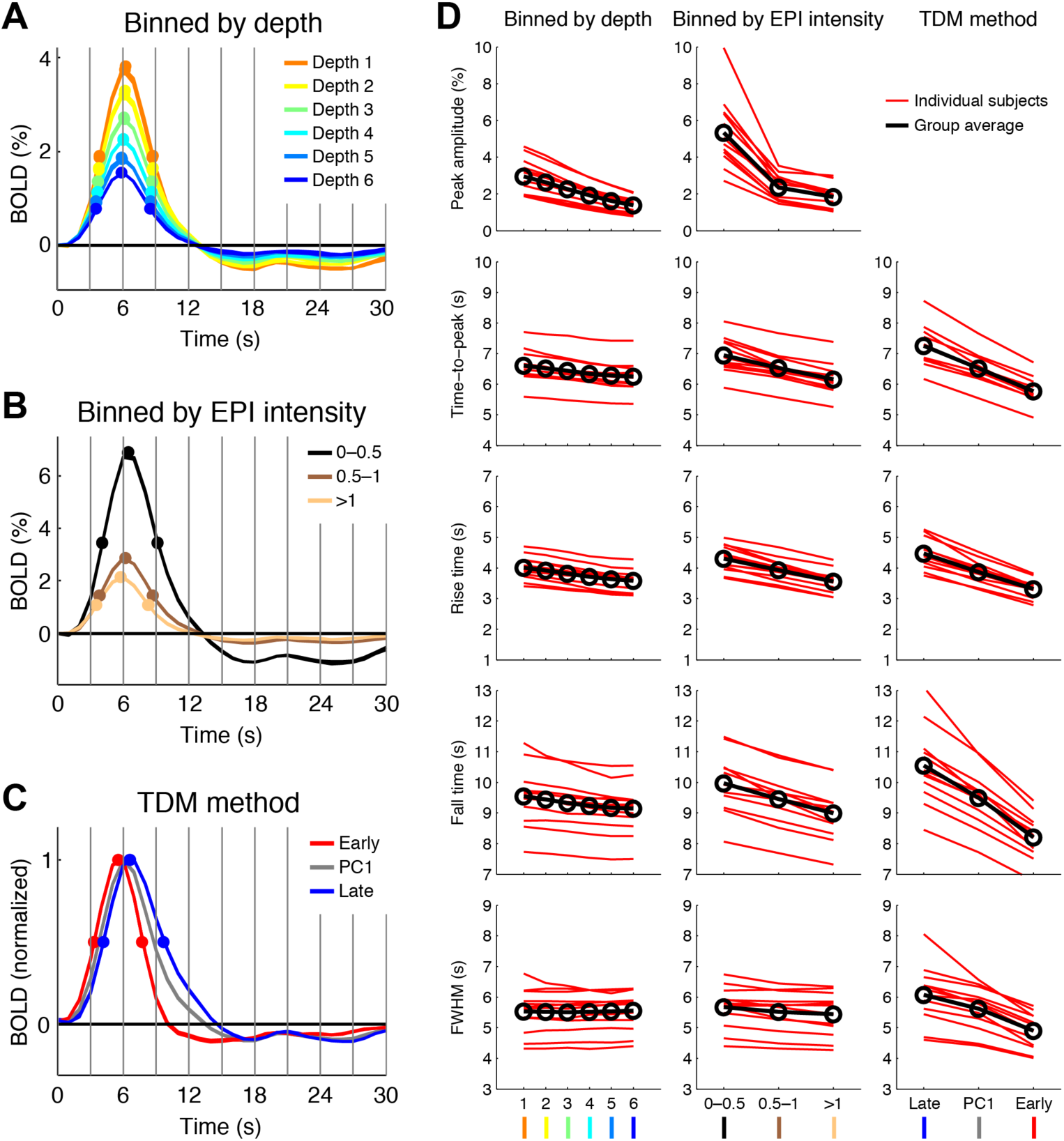
BOLD timecourses exhibit systematic variation in amplitude, delay, and width. Panels A–C show detailed results for Dataset D10 (detailed results for all datasets are shown in **Supplementary Figure 1**). *A*, Timecourses binned by cortical depth (Depth 1 is superficial; Depth 6 is deep). Timecourses from split-halves of the data are shown for each bin (the two sets of traces are nearly identical, indicating high reliability). Vertical gray lines mark 3-s intervals, a convention used throughout this paper. Solid dots mark timecourse peak, rise time (time at which the signal rises to half of the peak value), and fall time (time at which the signal falls to half of the peak value). Full-width-half-max (FWHM) is calculated as fall time minus rise time. *B*, Timecourses binned by EPI intensity. Same as panel A except binning is performed with respect to bias-corrected EPI intensity. *C*, Timecourses derived by TDM. The Early and Late timecourses derived by TDM are normalized to peak at 1. For comparison, we also show the first PC of the timecourses, also normalized to peak at 1. *D*, Summary of results across datasets. Timecourse metrics obtained for individual subjects (Datasets D1–D12) and corresponding group averages are shown.

**Figure 3.**
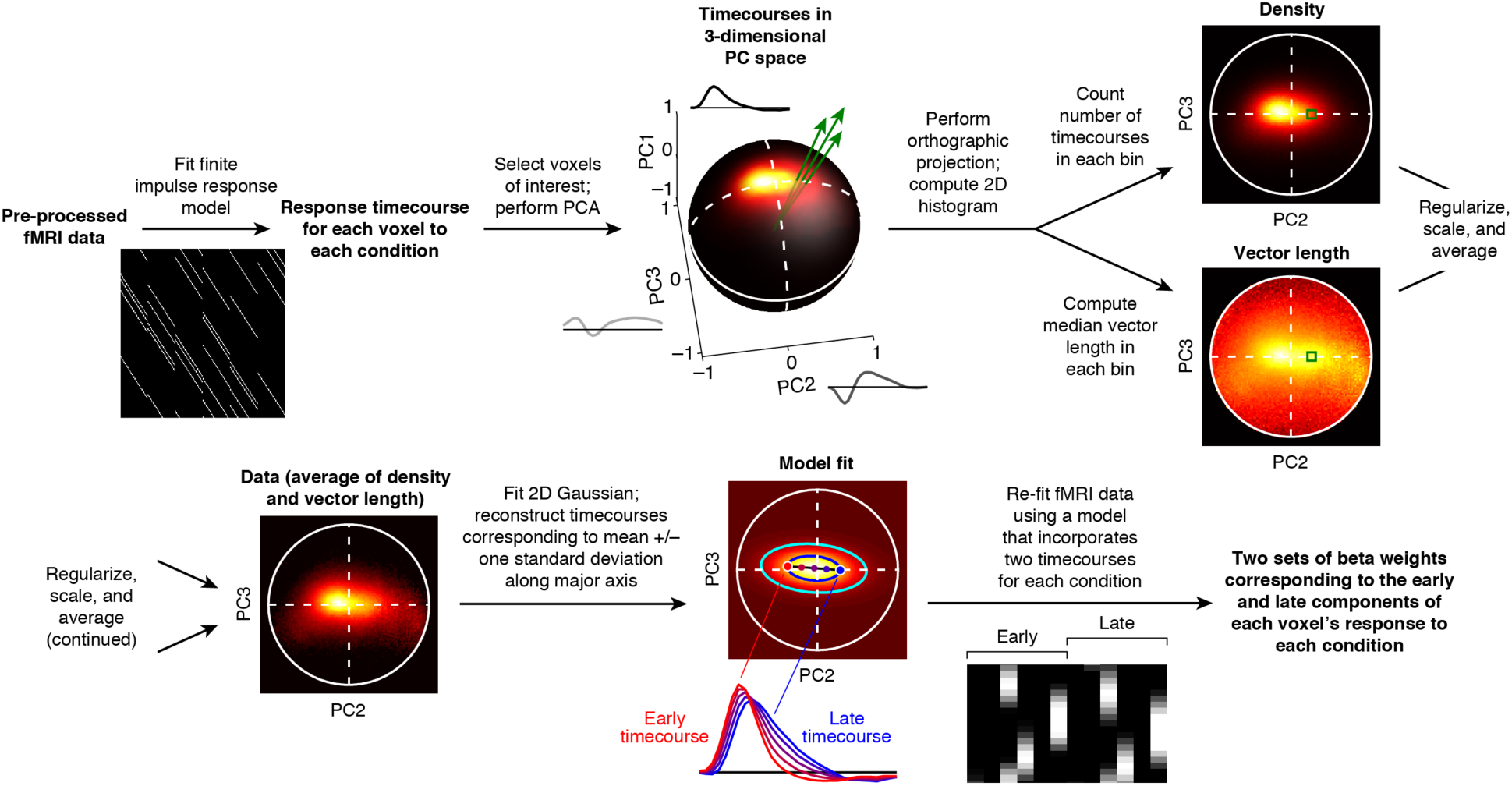
Schematic of the TDM method. Time-series data are fit with a finite impulse response model to estimate response timecourses. PCA is performed on the timecourses to reduce their dimensionality to 3. Using orthographic projection in the direction of the first PC, a 2D histogram image is calculated (*density*). The same projection and binning scheme is used to calculate an image representing timecourse amplitudes (*vector length*). The two images are combined and fit with a 2D Gaussian in order to determine an early timecourse and a late timecourse that together summarize the principal axis of variation. Finally, the time-series data are re-fit with a model incorporating the two timecourses.

**Figure 4.**
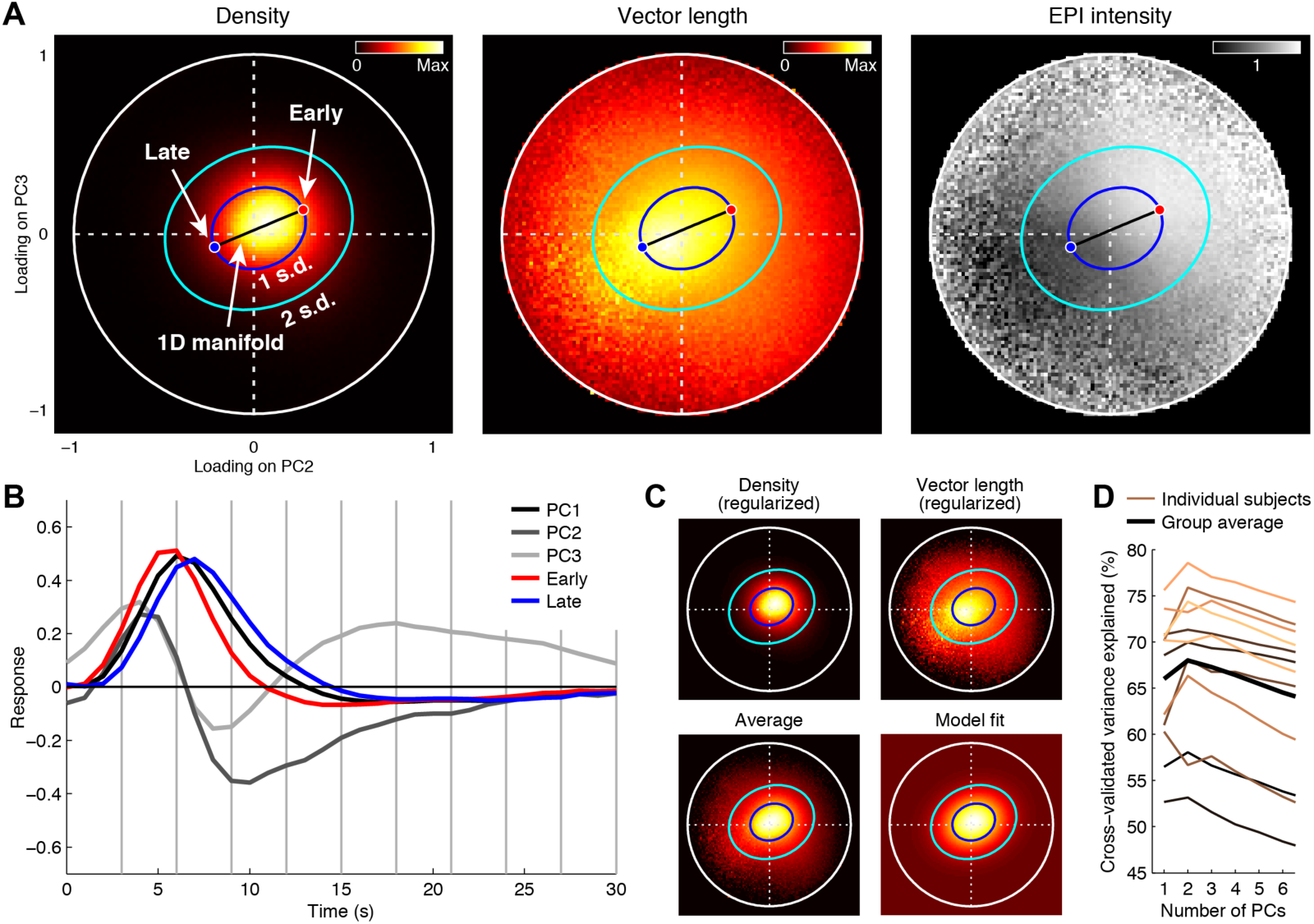
TDM captures timecourse variation along a 1D manifold. Panels A–C show results for Dataset D9 (results for all datasets are shown in **Supplementary Figures 2 and 3**). *A*, Quantities of interest. TDM calculates a density image indicating frequently occurring timecourse shapes (left), a vector-length image indicating timecourse amplitudes (middle), and an EPI-intensity image indicating bias-corrected EPI intensities (right). The black line indicates the identified 1D manifold that connects the Early and Late timecourses. (Ellipses, dots, and lines are identical in all plots.) *B*, Timecourses. All timecourses are unit-length vectors. *C*, Gaussian fitting procedure. Density and vector-length images are DC-subtracted, scaled, and truncated, producing regularized images (upper left, upper right). These images are then averaged (lower left) and fit with an oriented 2D Gaussian (lower right). *D*, Dimensionality of response timecourses. To confirm that three dimensions are sufficient to capture relevant variation in response timecourses, we perform PCA on split-halves of each dataset and assess how well a limited number of PC timecourses from one half reconstruct timecourses measured in the other half (i.e. cross-validation). Results for individual subjects (Datasets D1–D12) and the group average are shown.

**Figure 5.**
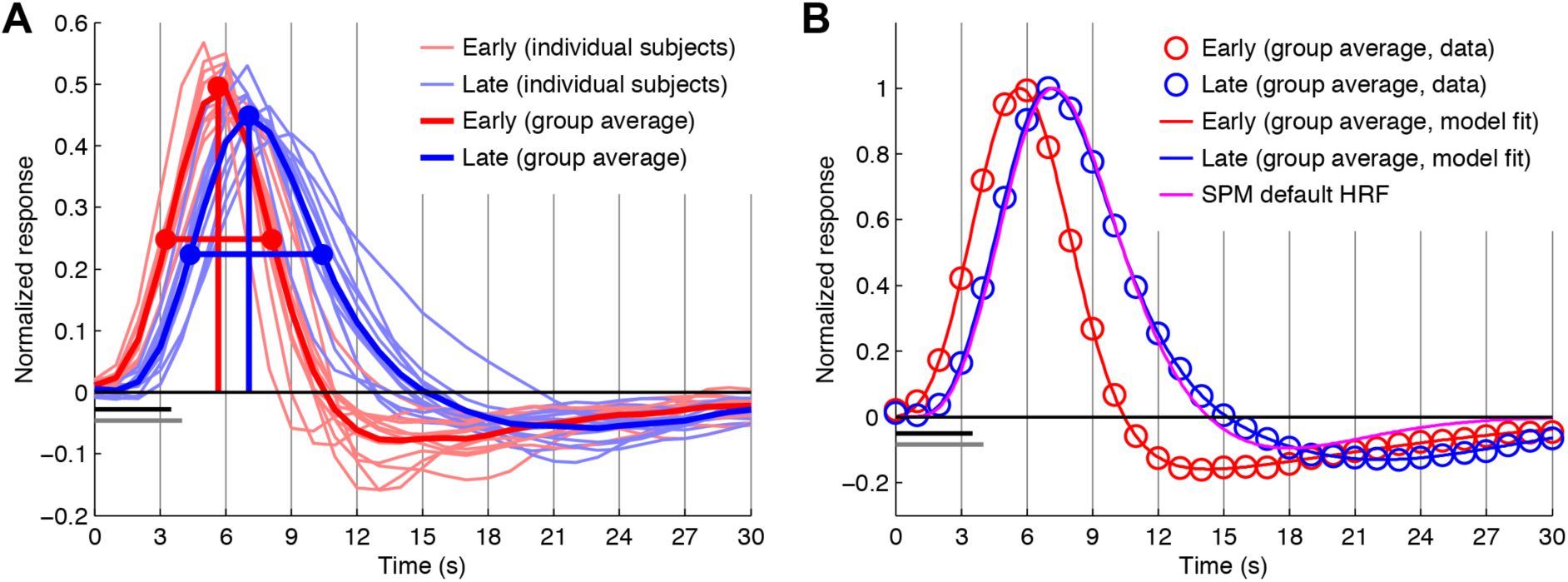
Early and late timecourses found by TDM. *A*, Summary of results. We plot unit-length-normalized Early and Late timecourses found in Datasets D1–D12 (thin lines) and their average (thick lines). Dots mark timecourse peaks and timecourse rise and fall times. Horizontal black and gray bars below the *x*-axis indicate the stimulus duration in the eccentricity and category experiments (3.5 s and 4 s, respectively). *B*, Group-average results and parametric model fit. Group-average Early and Late timecourses are normalized to peak at 1 (red and blue circles), and are fit with a double-gamma function as implemented in SPM’s *spm_hrf*.*m* (red and blue lines). The fitting was achieved by convolving a double-gamma function with a 4-s square wave and minimizing squared error with respect to the data. Estimated double-gamma parameters for the group-average Early and Late timecourses were [7.21 17.6 0.5 4.34 1.82 –3.09 50] and [5.76 21.6 1.11 1.72 3.34 0.193 50], respectively. As a comparison, we show the timecourse obtained using the default SPM parameters [6 16 1 1 6 0 32] (magenta line).

The second GLM model, termed Standard, is a GLM in which a canonical hemodynamic response function (*getcanonicalhrf*.*m*) is convolved with stimulus onsets to create a regressor for each experimental condition. We used six condition-splits, thereby producing six response estimates for each condition. Fitting the Standard model produced BOLD response amplitudes (‘betas’) with dimensionality *N* vertices × 6 depths × *M* conditions × 6 condition-splits.

The third GLM model, termed TDM, is a GLM in which two hemodynamic timecourses (Early, Late) are separately convolved with stimulus onsets to create two regressors for each experimental condition. The exact nature of these timecourses is determined by the TDM method as described in Section 2.8. We used six condition-splits, thereby producing six response estimates for each combination of condition and timecourse. Fitting the TDM model produced BOLD response amplitudes (‘betas’) with dimensionality *N* vertices × 6 depths × *M* conditions × 2 timecourses × 6 condition-splits.

GLMs were prepared and fit to the data using GLMdenoise (Charest et al., 2018; Kay et al., 2013). In GLMdenoise, the GLM consists of experimental regressors (which may take on different forms, as described above), polynomial regressors that characterize the baseline signal level in each run, and data-derived nuisance regressors. In the case of the FIR model, experimental regressors consisted of binary values (0s and 1s). In the case of the Standard and TDM models, experimental regressors consisted of the convolution of stimulus onsets (1s) with hemodynamic timecourses that are normalized to peak at 1 (for example, see **Figure 2C**). After fitting the GLMs, estimated betas were converted from raw scanner units to units of percent BOLD signal change by dividing by the mean signal intensity observed at each vertex and multiplying by 100.

Betas were further analyzed using simple summary metrics. To quantify overall BOLD activity at a given vertex, we calculated *mean absolute beta* (e.g. **Figure 6, left**) by averaging betas across condition-splits, taking the absolute value of the results, and then averaging across conditions. To summarize responses to the eccentricity stimuli, we calculated *peak eccentricity* (e.g. **Figure 6, middle**) by averaging betas across condition-splits, performing positive half-wave rectification (i.e. setting negative values to zero), and then calculating center-of-mass (Hansen et al., 2007). Specifically, center-of-mass was calculated as the weighted average of the integers 1–6 (corresponding to the 6 ring eccentricities from fovea to periphery) using the rectified betas as weights.

**Figure 6.**
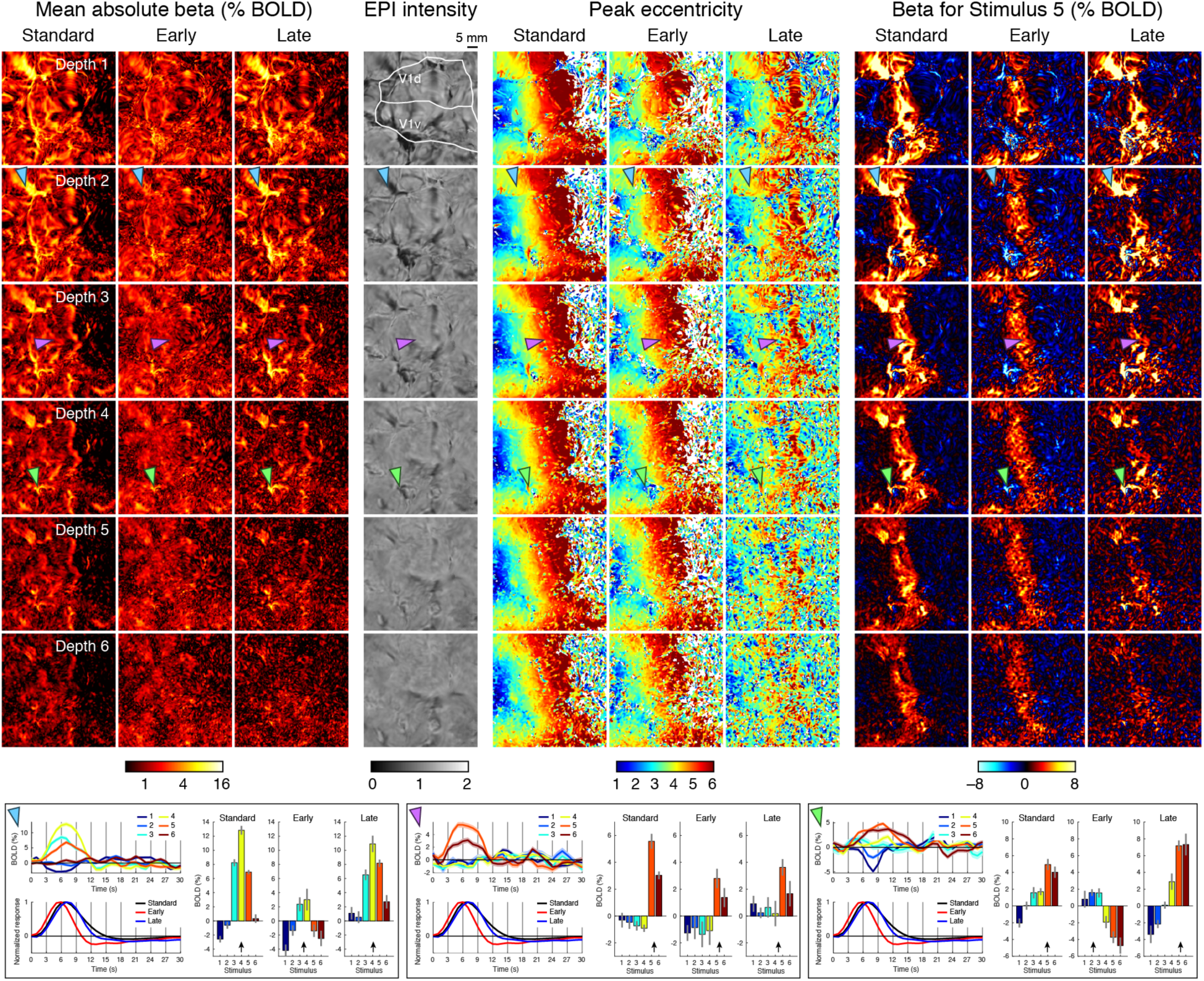
TDM decomposes brain activity patterns into early and late components. Here we show detailed results for Dataset D1; summary results for all datasets are provided in **Figure 7**. Rows correspond to different cortical depths for a small patch covering primary visual cortex (flattened surface, left hemisphere). Three versions of results are shown: the first (Standard) reflects betas from a GLM incorporating a single canonical HRF, while the next two (Early, Late) reflect betas from a GLM incorporating the timecourses found by TDM. Four types of images are shown from left to right: (1) absolute value of betas averaged across conditions, (2) bias-corrected EPI intensities, (3) peak eccentricity quantified as the center-of-mass of the six betas observed at each vertex, and (4) single-condition activity patterns for the fifth stimulus. Blue, purple, and green arrows mark vertices illustrated in greater detail in the inset boxes. Each inset box shows FIR timecourses (upper left) with ribbon width indicating standard error across condition-splits; canonical and TDM-derived timecourses (lower left); and the three versions of the betas (right) with error bars indicating standard error across condition-splits and black arrows indicating peak eccentricity. Notice that at the level of individual vertices, TDM decomposes BOLD responses into two sets of betas (Early, Late) that exhibit different stimulus selectivity.

### 2.7. Timecourse quantification and metrics

In some analyses (see **Figure 2A–B, Supplementary Figure 1**), we summarize the typical timecourse shape observed in a set of timecourses. This was accomplished using a PCA-based procedure (*derivehrf*.*m*). In the procedure, we first subtract off the mean of each timecourse. We then perform principal components analysis (PCA) on the entire set of timecourses and extract the first PC (this is the timecourse vector along which variance in the total set of timecourses is maximized). Next, we add a constant offset to the first PC such that the first time point equals 0, and flip the sign of the PC such that the mean over the range 0–10 s is positive. Finally, we calculate the weight that minimizes squared reconstruction error for each timecourse, compute the absolute value of these weights, and then scale the PC by the average weight. The motivation for de-meaning the timecourses prior to PCA is to suppress low-frequency noise present in weak BOLD responses. In general, the use of PCA for summarizing timecourse shape (Kay et al., 2008a) has advantages over simply computing the mean timecourse: PCA elegantly handles negative BOLD timecourses, and PCA allows timecourses with larger BOLD responses to have greater influence on the resulting timecourse shape (thereby producing more robust results).

Several timecourse metrics were computed (see **Figure 2, Figure 5, Supplementary Figure 1**). Given a timecourse, we upsampled the timecourse to a sampling rate of 0.01 s using sinc interpolation. We then identified the maximum of the resulting timecourse (*peak amplitude*) and its associated time (*time-to-peak*). We used linear interpolation to calculate the time at which the timecourse rises to half of the maximum value (*rise time*) and the time at which the timecourse falls to half of the maximum value (*fall time*). Finally, we computed the time elapsed between the rise time and the fall time (*full-width-at-half-max* or *FWHM*).

### 2.8. TDM method

#### 2.8.1. Theory

TDM is a data-driven technique that identifies a principal axis of timecourse variation present in a set of experimentally measured timecourses. It does this by examining timecourses projected into a low-dimensional space (3 dimensions) and extracting a 1-dimensional manifold (a line) that captures the variation of interest. The procedure can be viewed as a powerful method for summarizing and extracting the signal present in timecourses which considered individually (i.e. one response timecourse at a time) would likely be insufficiently reliable. In our fMRI measurements of responses to 4-s visual stimuli, we consistently find that one endpoint of the line corresponds to an early timecourse peaking at around 5–7 s and the other endpoint of the line corresponds to a late timecourse peaking at around 6–9 s (see **Figure 2D, Figure 5A, Supplementary Figure 1**). These timecourses are interpreted as reflecting hemodynamic responses from the microvasculature (e.g. capillaries and venules) and hemodynamic responses from the macrovasculature (e.g. veins), respectively. TDM then uses the identified timecourses in a regression model in order to decompose observed hemodynamic responses into early and late components. The researcher can choose to analyze further the early component, the late component, or both.

There are three main quantities involved in the TDM technique: *density*, referring to the timecourse shapes that tend to be present in the data; *vector length*, referring to the amplitudes of the timecourses in the data; and *EPI intensity*, referring to the bias-corrected EPI intensity of the vertex (or voxel) to which each timecourse belongs. TDM combines density and vector length and fits an oriented 2D Gaussian to the result in order to identify the 1-dimensional manifold. The motivation for incorporating vector length beyond density alone is to ensure that veins—which generate BOLD responses with large amplitudes but constitute only a fraction of the total set of responses—have sufficient influence on the determination of the manifold. Note that EPI intensity does not directly participate in the determination of the manifold, and can therefore provide useful validation of the manifold results (see **Figure 4A, Supplementary Figures 2–3**).

There are a few important conceptual points regarding the nature of the TDM method. For any given voxel (or vertex), the BOLD response to an experimental event is expected to reflect a mixture of early (microvasculature) and late (macrovasculature) timecourses. The specific proportions of these timecourses is expected to vary from voxel to voxel simply due to heterogeneity in the spatial structure of the vasculature (e.g., one voxel might be centered on a large vein, whereas another voxel may only partially overlap the vein). Different proportions of the timecourses manifest in TDM as different points along the 1-dimensional manifold, and these points collectively trace out an arc on the unit sphere (see **Figure 3**).

Another important point is that TDM is not equivalent to estimating a different hemodynamic response function for each voxel (e.g., Kay et al., 2008b, 2008a; Pedregosa et al., 2015). Using a single hemodynamic timecourse for different experimental conditions (time-condition separability) yields at most one amplitude estimate (beta) for each condition. In contrast, TDM allows experimental conditions to have different loadings on the early and late timecourses, and yields two amplitude estimates (betas) for each condition (see **Figure 6, inset boxes**). This is the critical feature that gives TDM the power to remove unwanted venous effects from a given voxel’s response.

Finally, note that the GLM analyses performed in TDM rely on the assumption of linear summation of BOLD responses over time. Our experiments, like many used in cognitive neuroscience, involve a large number of trials that are presented fairly rapidly in order to maximize statistical power (e.g. less than 10 s of rest in between trials). In such experiments, it is a practical necessity to assume temporal linearity. Moreover, nonlinear effects are likely to average out when using randomized experimental designs. Nevertheless, the accuracy with which TDM identifies timecourses may be limited if there exist nonlinear effects (Friston et al., 1998b) and especially if these nonlinearities vary for different types of vasculature (Goodyear and Menon, 2001; Thompson et al., 2014; Zhang et al., 2009).

#### 2.8.2. Algorithm

The TDM algorithm starts with a set of timecourses and determines a pair of timecourses that characterize the overall variation in the timecourses. The following are the steps in the TDM algorithm (*extracthrfmanifold*.*m*):

1. *Perform PCA on the timecourses*. We collect timecourses into a 2D matrix of dimensionality *L* timecourses × *T* time points, and then perform singular value decomposition. This produces a matrix with dimensionality *T* time points × *T* eigenvectors where columns correspond to principal component (PC) timecourses in decreasing order of variance explained. We flip the sign of the first PC if necessary to ensure that the mean of the timecourse is positive over the range 0–10 s. Note that previous studies have applied PCA to fMRI response timecourses but in different contexts (d’Avossa et al., 2003; Woolrich et al., 2004).
2. *Use PC1–PC3 to define a 3-dimensional space for further analysis*. Our convention for visualization is that PC1 points out of the page (positive *z*-axis), PC2 points to the right (positive *x*-axis), and PC3 points to the top (positive *y*-axis).
3. *Map timecourses onto the unit sphere*. We project each timecourse onto PC1, PC2, and PC3, and mirror coordinates across the origin if necessary to ensure that the loading on PC1 is positive. (Thus, negative BOLD timecourses are flipped and treated in the same way as positive BOLD timecourses.) Each timecourse is represented by a set of coordinates (loadings) and can be interpreted as a 3-dimensional vector. We normalize each vector to unit length (thereby placing the vector on the unit sphere), and also save the original vector length for later use.
4. *Calculate a 2D image that represents density*. We orthographically project timecourses onto the *xy* plane, and then calculate a 2D histogram. This produces a 2D image where pixel values represent frequency counts. This image indicates typical timecourse shapes found in the data.
5. *Calculate 2D images that represent vector length and EPI intensity*. Using the same orthographic projection and binning scheme of Step 4, we calculate the median vector length of the timecourses found in each bin. We also calculate the median bias-corrected EPI intensity of the voxels (vertices) associated with the timecourses in each bin. This produces 2D images where pixel values represent vector lengths (indicating timecourse amplitudes) and EPI intensities (indicating static susceptibility effects caused by veins), respectively.
6. *Regularize the density image by subtracting a DC bias*. We distribute a collection of particles on the unit sphere (S2 Sampling Toolbox, https://www.github.com/AntonSemechko/S2-Sampling-Toolbox), assign timecourses to their nearest particles, and count the number of timecourses associated with each particle. We then calculate a histogram of these counts and determine the bin *B* with the highest frequency. Finally, we stochastically subsample the timecourses such that the number of timecourses associated with each particle is reduced by the middle value of bin *B*. The resulting subsampled timecourses are used to generate a new density image.
7. *Scale the density image*. The regularized density image from Step 6 is scaled such that 0 maps to 0 and the maximum value maps to 1. Values are then truncated to the range [0,1].
8. *Regularize the vector-length image by subtracting a DC bias*. This is accomplished in a similar manner as Step 6: we calculate a histogram of the values in the vector-length image, determine the bin *B* with the highest frequency, and subtract the middle value of bin *B* from all image pixels.
9. *Scale the vector-length image*. The regularized vector-length image from Step 8 is scaled such that 0 maps to 0 and the maximum value maps to 1. Values are then truncated to the range [0,1].
10. *Average the density and vector-length images*. Although the default behavior is to simply average the density and vector-length images, if the user desires a different weighting (e.g. giving more weight to the vector-length image), a flag can be used (*opt*.*vlengthweight*).
11. *Fit 2D Gaussian*. The image resulting from Step 10 is fit with a 2D Gaussian. The Gaussian is controlled by two parameters specifying the center, two parameters specifying the spreads along the major and minor axes, a rotation parameter, a gain parameter, and an offset parameter. In model fitting, the error metric is set up such that the image is interpreted as a probability distribution (pixels with larger values reflect higher density and thus contribute more heavily to the error metric).
12. *Extract two points along the major axis of the Gaussian*. We determine points corresponding to the mean ± one standard deviation along the major axis of the Gaussian. The choice of one standard deviation is somewhat arbitrary but appears to produce satisfactory results.
13. *Reconstruct timecourses corresponding to the identified points*. We place the points determined in step 12 on the unit sphere, and use their associated coordinates to weight and sum the PC1, PC2, and PC3 timecourses. This yields two reconstructed timecourses. Based on time-to-peak (see Section 2.7), we label one timecourse as ‘Early’ and the other timecourse as ‘Late’.

#### 2.8.3. Application of algorithm

We obtained timecourses by fitting an FIR model to the fMRI time-series data (as described in Section 2.6). We then calculated the amount of variance explained (*R*^2^) by the FIR model. In order to focus the TDM algorithm on cortical locations with BOLD responses, we selected all vertices within a region of interest (see Section 2.9) that exceeded an automatically determined threshold (specifically, the value at which the posterior probability switches between two Gaussian distributions fitted as a mixture model to the data; see *findtailthreshold*.*m*). This produced a set of timecourses with dimensionality *P* vertices × *M* conditions × 31 time points × 2 condition-splits. We applied the TDM algorithm to the timecourses averaged across the condition-splits (thus, reflecting the entire dataset) and also to the timecourses from each condition-split separately in order to assess reliability. In both cases, the number of timecourses given to the TDM algorithm is *L* = *P*\**M* and the number of time points is *T* = 31. After completion of the TDM algorithm, the identified timecourses (Early, Late) were incorporated into a GLM model to decompose the fMRI time-series data into early and late components (see Section 2.6).

#### 2.8.4. Alternative ICA-based procedure

Independent components analysis (ICA) is a potential alternative method for determining latent timecourses. We designed an ICA-based procedure (*icadecomposehrf*.*m*) that serves as a drop-in replacement for the TDM algorithm described above. In the procedure, we take the two condition-split versions of the FIR-derived timecourses and perform ICA on each set of timecourses (FastICA Toolbox, https://research.ics.aalto.fi/ica/fastica/, default nonlinearity). This produces 31 independent component (IC) timecourses for each split of the data. Because ICA does not provide a natural ordering or grouping of the ICs, the challenge is to determine which specific pair of ICs to use as the latent timecourses in TDM.

To determine the pair of ICs to use, we devised the following heuristic procedure. We first greedily order the ICs within each split to maximize variance explained. That is, we choose the IC that maximizes variance explained in the timecourses, choose a second IC that, when combined with the first, maximizes variance explained, and so on. For normalization purposes, we flip each IC if necessary to ensure that it is positive over the range 0–10 s and normalize it to unit length. Next, we perform greedy matching in order to match the ICs in the second split of the data to the ICs in the first split. Specifically, we choose the IC in the second split that is most similar in a squared-error sense to the first IC in the first split, choose the remaining IC in the second split that is most similar to the second IC in the first split, and so on. To reduce the number of ICs under consideration, we select ICs that both exhibit high consistency across the two splits of the data (*R*^2^ > 50%) and are within the top 50% of the ICs in the first split (with respect to the ordering based on variance explained). Finally, from the ICs that meet these selection criteria, we perform an exhaustive search to determine the unique pair of ICs that explain the most variance in the original timecourses. Based on time-to-peak (see Section 2.7), we label one IC as ‘Early’ and one IC as ‘Late’.

### 2.9. Region-of-interest (ROI) definition

We used two regions-of-interest (ROIs). For the eccentricity experiment, we used the union of visual areas V1, V2, and V3 from a publicly available atlas of visual topography (Wang et al., 2015). For the category experiment, we used a manually defined region in occipital, parietal, and temporal cortex that covers visually responsive vertices (same region used in Kay et al., 2019). Both ROIs were defined in FreeSurfer’s *fsaverage* space and backprojected to individual subjects for the purposes of vertex selection.

## 3. Results

We collected BOLD fMRI measurements in human visual cortex while subjects viewed briefly presented visual stimuli (3.5-s or 4-s duration). The main datasets are D1–D5 in which a 7T gradient-echo 0.8-mm 2.2-s protocol was used to measure responses to rings at different eccentricities (**Figure 1A**). Additional datasets D6–D12 used the same acquisition protocol but involved images of different semantic categories (e.g. faces). The data were pre-processed by performing one temporal interpolation that corrected slice-time differences and prepared the data at a 1-s sampling rate (**Figure 1B**) and one spatial interpolation that corrected head motion and EPI distortion and sampled the fMRI volumes at the locations of cortical surface vertices. Given the high resolution of the fMRI data, we prepared multiple cortical surface representations at different depths through the gray matter.

### 3.1. Systematic variation in timecourse amplitude, delay, and width

To investigate BOLD timecourse characteristics, we analyzed the time-series data in each dataset using a finite impulse response model (0–30 s, 31 time points). This produced, for each surface vertex, an estimate of the BOLD response timecourse to each experimental condition. We then binned these timecourses either with respect to cortical depth or with respect to EPI intensity. For each bin, we summarized the timecourses found in that bin using a PCA-based procedure (see Methods for details).

Results for a representative dataset are shown in **Figures 2A–B**. Proceeding from inner cortical depths (Depth 6) to outer cortical depths (Depth 1), we observe an increase in timecourse amplitude and an increase in timecourse delay (**Figure 2A**). Proceeding from darker EPI intensities (0–0.5) to lighter EPI intensities (>1), we again observe increases in amplitude and delay but also a small increase in timecourse width (**Figure 2B**). Summarizing results across different datasets (reflecting different subjects and experiments), we see that these effects are consistently observed (**Figure 2D**). Notice that although different datasets show similar patterns in relative timing (e.g., time-to-peak is longer for low EPI intensity than for high EPI intensity), there is large variance in absolute timing across datasets (e.g., time-to-peak is approximately 8 s in one dataset but 6 s in another dataset). This is consistent with well-established observations of variability of hemodynamic response functions across subjects (Handwerker et al., 2012).

Notice that the variations in amplitude (**Figure 2D**, top row) and the variations in timecourse delays (**Figure 2D**, middle rows) are more pronounced when binning by EPI intensity (**Figure 2D**, middle column) than when binning by cortical depth (**Figure 2D**, left column). We suggest that the underlying source of these effects consists in the high-amplitude, delayed BOLD responses carried by macroscopic veins. Binning by EPI intensity provides a relatively direct proxy for where these venous effects occur (Kay et al., 2019; Menon et al., 1993; Ogawa et al., 1990). In contrast, binning by cortical depth provides a less direct proxy for these effects (e.g., pial veins affect outer cortical depths more than inner cortical depths). Thus, binning by EPI intensity will tend to accentuate and highlight the amplitude and delay effects.

The fact that veins carry delayed BOLD responses has been previously shown (de Zwart et al., 2005; Kim and Ress, 2017; Lee et al., 1995; Siero et al., 2011). Prior studies have also demonstrated increased temporal delays at superficial cortical depths (Kim and Ress, 2017; Siero et al., 2011). Thus, the observations we make here are not novel, but provide reassurance that these effects can be reproduced in our dataset and establish a starting point for the development of the TDM method.

### 3.2. TDM provides a method for visualizing timecourse variation

As we have seen, there is systematic variation in the hemodynamic timecourses observed in our data. Given that capturing timecourse variation lies at the core of the TDM method, we are now ready to examine results of applying TDM to our data.

The first step in TDM is to visualize distributions of response timecourses in a low-dimensional space (**Figure 3**). Specifically, we use principal components analysis (PCA) to determine the three orthogonal timecourses that account for the most variance in the given timecourses. We then use these three timecourses as axes of a 3D space (PC1, PC2, PC3) in which each of the original timecourses corresponds to one point, or vector, in this space. To visualize the results, we map the timecourse vectors to the unit sphere (by normalizing them to unit length) and use an orthographic projection to visualize the density of the timecourses. Such a visualization reveals typically occurring timecourse shapes, independent of timecourse amplitude (see ‘Density’ image in **Figure 3**). We separately visualize the amplitude of the timecourses by computing the length of the original timecourse vectors and repeating the orthographic visualization (see ‘Vector length’ image in **Figure 3**).

Applying these visualization procedures to a representative dataset (similar results are obtained in every dataset; see **Supplementary Figures 2–3**), we find that timecourse shapes typically reside near the pole of the unit circle where PC1 is maximal, with some variability around this pole (**Figure 4A, left**). We find that timecourse amplitudes are large in a similar portion of the space except for a small extension towards the lower left (**Figure 4A, middle**). A separate plot shows the actual timecourses associated with the three PCs that define the axes of the space (**Figure 4B**). This plot shows that timecourse shapes generally resemble a canonical hemodynamic response timecourse (**Figure 4B, black line**), with a major axis of variation corresponding to different loadings on a timecourse that shifts the peak of the canonical timecourse either earlier or later in time (**Figure 4B, dark gray line**). Note that the general shapes of the PC timecourses are similar to those found in other applications of PCA to fMRI timecourses (d’Avossa et al., 2003; Woolrich et al., 2004).

We emphasize the usefulness of the visualizations performed in TDM: the visualizations summarize large amounts of data (response timecourses) in a manner that highlights robust signals and clearly delineate effects of interest in the data. We suggest that the visualizations may be generally useful for investigating timecourse variations across voxels, brain areas, and/or subjects. Although the use of three dimensions in the visualizations is appealing because it is practical to create pictorial representations of a small number of dimensions, visualizing timecourses in only three dimensions might provide an incomplete characterization of the full diversity of timecourses. To investigate this issue, we performed a cross-validation analysis to determine the number of timecourse dimensions necessary to capture signals of interest in the response timecourses (**Figure 4D**). We find that in every dataset, cross-validation performance is maximized using three or fewer PCs. Thus, it appears that using three PC dimensions is sufficient and that additional dimensions would likely be dominated by measurement noise.

### 3.3. TDM identifies an axis of timecourse variation

The second step in TDM is to identify an axis that captures the major variation in the observed response timecourses. As seen in the earlier visualizations (**Figure 4A**), response timecourses empirically occupy a small portion of the 3D space (i.e., timecourse density and vector length are confined to a small section of the unit sphere). Furthermore, the timecourses can be approximated in the 3D space by a simple line segment (arc) that describes variation with respect to timecourse delay, i.e. early vs. late. The interpretation that we adopt here is that (i) the endpoints of the line segment correspond to latent hemodynamic timecourses associated with the microvasculature and the macrovasculature and (ii) any single observed timecourse is simply a mixture of these two latent timecourses plus measurement noise (which causes deviation away from the line). There are a variety of reasons why different mixtures might be observed across voxels in a given dataset. One simple reason is heterogeneity in the spatial structure of the vasculature: some voxels might be in close proximity to a large vein whereas other voxels may largely comprise capillaries and small venules. (Note that one might have expected to find discrete clusters or modes in the timecourse distribution, but this does not appear to be the case in our measurements. It is possible that clustering might be observed at higher spatial resolutions.)

To calculate the axis of variation, TDM combines density and vector length (**Figure 3, lower left**), fits a 2D Gaussian to the result (**Figure 3, bottom**), and extracts points positioned at plus and minus one standard deviation along the major axis of the fitted Gaussian (**Figure 3, bottom, red and blue points**). Examining results obtained on the representative dataset, we see that the procedure works well: the combined image resembles both density and vector length (**Figure 4C, bottom left**), the fitted Gaussian approximates the data (**Figure 4C, bottom right**), and the extracted points reside in sensible locations (**Figure 4A, first two plots**). We reconstruct timecourses corresponding to the two extracted points (**Figure 4B, red and blue lines**), and then label the timecourses ‘Early’ and ‘Late’ based on the time-to-peak of the reconstructed timecourses. Note that different mixtures of the Early and Late timecourses trace out an arc (line) on the unit sphere (**Figure 4A, black line**) and result in a continuum of timecourse shapes (**Figure 3, bottom**). This arc is the 1D manifold that describes timecourse variation in the dataset.

We interpret the Early and Late timecourses as reflecting the microvasculature (e.g. capillaries and venules) and macrovasculature (e.g. veins), respectively. Does this interpretation have face validity? We offer several lines of reasoning that suggest validity. First, we find that the Late timecourse is consistently associated with large vector length (see **Figure 4A, middle plot** and **Supplementary Figure 2**), indicating that response timecourses resembling the Late timecourse tend to have large BOLD amplitudes. This makes sense given that veins exhibit large percent BOLD signal changes (Lee et al., 1995; Menon et al., 1993). Second, if we use the same visualization methods (orthographic projection of the unit sphere) to examine the relationship between timecourse shape and bias-corrected EPI intensity, we find that the Late timecourse is consistently positioned in a zone of low EPI intensity (see **Figure 4A, right plot** and **Supplementary Figure 2**). Since the TDM procedure does not make use of EPI intensities, this is an empirical finding that provides further evidence of validity, as it is known that veins cause static susceptibility effects in EPI images (Kay et al., 2019; Menon et al., 1993; Ogawa et al., 1990). Third, the idea that veins exhibit delayed BOLD responses is consistent with several previous experimental studies (de Zwart et al., 2005; Kim and Ress, 2017; Lee et al., 1995; Siero et al., 2011). Finally, a biophysical model of vascular dynamics has been proposed, and this model provides a potential explanation for why temporal delays occur in veins (see Fig. 6 in Havlicek and Uludağ, 2020).

As an additional sanity check, let us compare the Early and Late timecourses identified by TDM against the timecourse inspections presented earlier. We see that the Early and Late timecourses (**Figure 2C**) resemble the ones found through binning by cortical depth (**Figure 2A**) and binning by EPI intensity (**Figure 2B**), but are somewhat more extreme in nature. For example, time-to-peak is earlier for the Early timecourse and later for the Late timecourse compared to the binning-based timecourses. These effects can be seen more clearly in the quantitative summary (**Figure 2D, right column**). Thus, TDM appears to be extracting sensible timecourses (see also **Supplementary Figure 1**). Finally, independent components analysis (ICA) is a commonly used statistical method for deriving latent structure in a set of data, and is commonly applied to fMRI data. We show that applying an ICA-based procedure yields results similar to those obtained from TDM in some datasets, but divergent and unsatisfactory results in other datasets (**Supplementary Figure 2**). We provide additional thoughts regarding ICA in the Discussion.

### 3.4. Summary of TDM timecourses across subjects

For a comprehensive assessment of the TDM results, we plot the TDM-derived Early and Late timecourses obtained for each dataset (**Figure 5A, thin lines**). We see that on the whole, the timecourses are stereotyped and largely consistent across datasets. Keep in mind that TDM is a completely data-driven technique: it makes no assumptions about timecourse shapes that are extracted from the data, and assumes only that timecourses can be characterized in a low-dimensional subspace and can be summarized by a one-dimensional manifold. Thus, the fact that the derived timecourses resemble typical hemodynamic response functions is an empirical finding, and indicates that the TDM technique successfully derives reasonable timecourses when applied to individual datasets.

While there are qualitative similarities of the Early and Late timecourses across datasets, there is also substantial quantitative variability, consistent with the well-established observation that BOLD timecourses exhibit variability across subjects (Handwerker et al., 2012). It is important to note that the observed variability in timecourses is not due to measurement noise: split-half analyses show that the TDM-derived timecourses are highly stable across splits of each dataset (**Supplementary Figure 1, right column**). These results suggest that in order to achieve the most accurate characterization of responses, one should tailor timecourses to what is empirically observed in individual datasets.

To summarize overall timecourse results, we compute the group average for the Early and Late timecourses (**Figure 5A, thick lines**). We see that these timecourses are similar in overall shape and differ primarily in their delay and width. We furthermore fit each group-average timecourse using a double-gamma function, and find that smooth parametric functions characterize the empirical results quite well (**Figure 5B**). Finally, as a point of comparison, we plot the predicted timecourse for a 4-s event using the default double-gamma hemodynamic response function implemented in SPM. Interestingly, this timecourse (**Figure 5B, magenta line**) coincides extremely well with the group-average Late timecourse (**Figure 5B, blue line**). This makes sense, given that the default timecourse parameters in SPM were derived from fMRI measurements conducted at low (2T) magnetic field strength (Friston et al., 1998a) where the BOLD response is dominated by contributions from large vessels (Haacke et al., 1994; Uludağ et al., 2009).

### 3.5. Decomposition using TDM timecourses removes artifacts from cortical maps

To summarize thus far, we have used TDM to derive Early and Late timecourses in each dataset, and we interpret these as reflecting responses from the microvasculature and macrovasculature, respectively. The final step of TDM involves re-analyzing the time-series data using a GLM that incorporates the Early and Late timecourses time-locked to the onset of each experimental condition. Fitting this GLM produces, for each vertex (or voxel) and condition, an estimate of the BOLD response amplitude from the microvasculature and an estimate of the BOLD response amplitude from the macrovasculature. These response amplitudes, or betas, can then be used in subsequent analyses according to the goals of the researcher; for example, one can choose to ignore the beta associated with the Late timecourse and focus on the beta associated with the Early timecourse. Note that the Early and Late timecourses are not necessarily orthogonal and are in fact often quite correlated (in **Figure 4A**, notice the close proximity of the Early and Late timecourses on the unit sphere). Thus, in the GLM model, the two timecourses are, in a sense, competing to account for the response timecourses observed in the data.

To assess the quality of the Early and Late betas, we generate cortical surface visualizations and compare these against visualizations of betas obtained using a standard GLM that incorporates a single canonical hemodynamic response function time-locked to each condition. We focus specifically on visualizations for the main datasets D1–D5 which involved presentation of stimuli that vary in eccentricity. This is because studies of the visual system provide well-established ‘ground truth’ expectations for neural activity patterns elicited by stimuli varying in eccentricity: in brief, neurons in early visual cortex respond selectively to stimuli at specific eccentricities, and the preferred eccentricity varies smoothly from foveal to peripheral eccentricities along the posterior-to-anterior direction (Wandell and Winawer, 2011).

Inspecting results for a representative dataset (**Figure 6**; other datasets shown in **Figure 7**), we first consider the overall magnitudes of the betas obtained with the different GLMs. The Standard betas appear to be roughly the sum of the Early and Late betas. We find that the Early and Late betas are both relatively large in magnitude (**Figure 6, first three columns**), indicating that both Early and Late timecourses contribute substantially to measured BOLD responses. However, the Early and Late betas show strikingly different patterns. The Early betas are relatively homogeneous across the cortical surface, relatively flat across cortical depth, and are moderate in size at around 1–4% signal change. In contrast, the Late betas are quite heterogeneous across the cortical surface (sparsely distributed), heavily biased towards outer cortical depths, and are sometimes quite large in size, reaching 10% or more signal change. The observation of additional activations arising in the Late betas is consistent with the fact that the BOLD point-spread function appears larger when sampling late in the BOLD response (Shmuel et al., 2007). Furthermore, comparing the spatial pattern of the Late betas against the spatial pattern of bias-corrected EPI intensities (**Figure 6, fourth column**), we see a general correspondence between the sparsely distributed locations where very large BOLD responses are observed and regions with dark EPI intensity. This co-localization of Late betas and dark EPI intensity is consistent with the earlier observation that timecourse shapes resembling the Late timecourse originate from vertices with dark EPI intensity (see **Figure 4A, right column**). These several observations (sparse distribution, outer depth bias, large BOLD signal change, dark EPI intensity) strongly suggest that the Late betas reflect responses from the macrovasculature.

**Figure 7.**
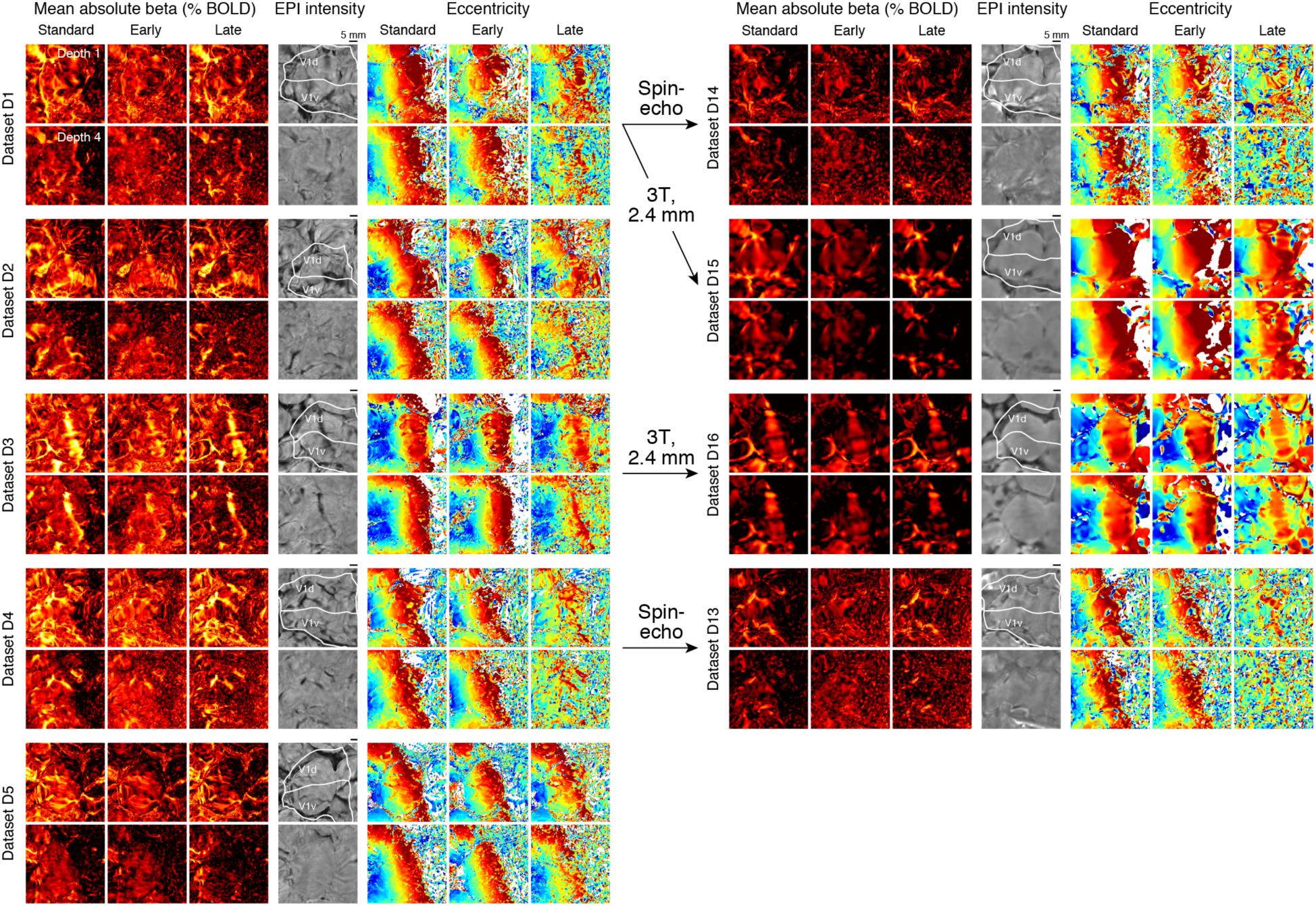
Decomposition of brain activity patterns across datasets and acquisition protocols. Same format as Figure 6, except only two cortical depths (Depths 1 and 4) are displayed. On the left are results obtained using high-resolution (0.8-mm) 7T gradient-echo (Datasets D1–D5). On the right are results obtained using high-resolution (1.05-mm) 7T spin-echo (Datasets D13–D14) and low-resolution (2.4-mm) 3T gradient-echo (Dataset D15–D16). These alternative acquisition protocols were conducted in the same subjects as the high-resolution gradient-echo protocol (correspondence indicated by arrows).

We next consider maps of peak eccentricity tuning, calculated as the center-of-mass of the betas corresponding to different stimulus eccentricities (**Figure 6, fifth through seventh columns**). All three versions of the betas (Standard, Early, Late) exhibit the expected smooth large-scale progression from foveal (blue) to peripheral (red) eccentricities as one moves posterior (left) to anterior (right) in early visual cortex. However, the quality or robustness of the eccentricity map is highest for the Standard betas, moderately high for the Early betas, and relatively low for the Late betas. Moreover, for the Late betas, there is a substantial decrease in quality moving from outer to inner cortical depths; this is consistent with the sharp fall-off in the magnitude of betas moving from outer to inner depths, as seen previously (see **Figure 6, third column**). We also observe that although large-scale eccentricity patterns are similar across the three versions of the betas, the maps show divergence at a finer scale. In particular, there are artifacts in eccentricity tuning that are present in the Standard and Late betas but absent in the Early betas. We illustrate a few example vertices in further detail (**Figure 6, arrows and inset boxes**); in these vertices, the tuning derived from the Early betas is better matched to expectations (based on inspection of neighboring sections of cortex and the large-scale eccentricity map). This is an important outcome and suggests that the Early betas isolate a component of the data that is more closely matched to responses from the microvasculature and avoids unwanted responses from the macrovasculature.

Finally, for further clarification of these results, we examine activity patterns elicited by a single experimental condition (**Figure 6, eighth through tenth columns**). Based on known tuning properties of early visual cortex (Wandell and Winawer, 2015, 2011), we expect a relatively compact ‘stripe’ of positive activity extending along the superior-inferior direction in cortex. All versions of the activity pattern (Standard, Early, Late) indeed show evidence of a stripe. However, there are differences, recapitulating effects observed earlier. Specifically, the Early beta is relatively homogeneous within the extent of the stripe, relatively flat across cortical depth, and moderate in size within the stripe at around 4% signal change. In contrast, the Late beta is heterogeneous across the cortical surface, heavily biased to outer cortical depths, and often very large in size, exceeding 8% signal change. These results demonstrate that TDM is capable of decomposing single-condition activity patterns into Early and Late components, with the former component more closely matching ground-truth expectations. Note that the ground-truth expectation is not necessarily a perfectly flat depth profile. This is because vascular density varies, to some degree, across cortical depth (Schmid et al., 2019). Moreover, our 0.8-mm measurements sampled within gray matter are certainly susceptible to partial volume effects from cerebrospinal fluid and white matter.

### 3.6. Quantification of TDM improvements

Although surface visualizations convey a large amount of information, they are qualitative. We therefore calculated quantitative metrics to more rigorously assess the results. First, we constructed histograms of the distributions of betas obtained under the different GLMs (**Figure 8A**). These distributions have long tails (**Figure 8A, bottom**) that appear to correspond to very large BOLD responses from the macrovasculature (see **Figure 6**). We quantified the magnitude of these tails using kurtosis, a metric that is large for heavy-tailed distributions. We find that Late betas have very high kurtosis, whereas Early betas have relatively low kurtosis (**Figure 8B**). Standard betas have an intermediate level of kurtosis, consistent with the interpretation that Standard betas behave essentially like an average or mixture of the Early and Late betas. In addition to quantifying the tails of the distributions, we quantified the overall magnitude of the betas by calculating the standard deviation of each distribution. We find that unlike kurtosis, the standard deviations of the beta distributions are similar across the three versions of the betas (**Figure 8C**). These results confirm that Early and Late betas are generally large in magnitude (indicating that BOLD responses are observed for both Early and Late timecourses), but Late betas have the unique feature of sometimes reaching extremely high values. This is consistent with the interpretation of Late betas as reflecting macrovascular responses.

**Figure 8.**
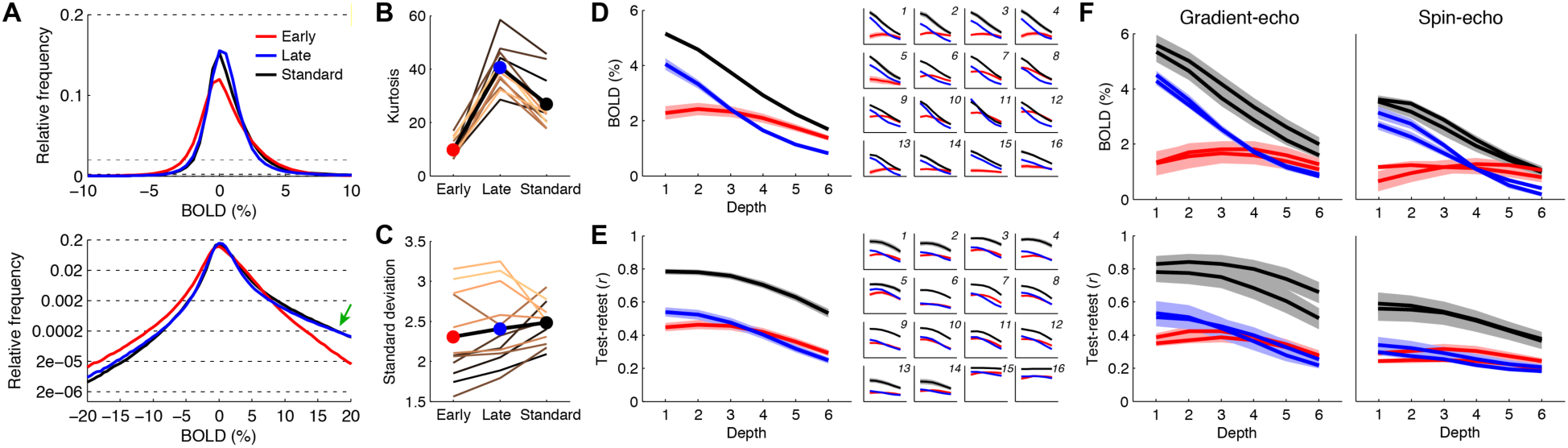
Quantitative assessment of BOLD amplitude estimates provided by TDM. *A*, Histogram of BOLD amplitudes. The top plot shows distributions of BOLD amplitudes aggregated across Datasets D1–D12; the bottom plot shows the same results on a log scale and with a wider *x*-axis range. *B*, Kurtosis of BOLD amplitudes. Results are shown for individual datasets (thin lines) and the group average (thick black line). *C*, Standard deviation of BOLD amplitudes. Same format as panel B. *D*, Cortical depth profiles of BOLD amplitudes. The main plot shows the average depth profile observed in Datasets D1–D12, with ribbons indicating standard error across datasets; the inset plots show results for individual datasets (D1–D16), with ribbons indicating standard error across conditions (same axis limits as the main plot). *E*, Test-retest reliability of BOLD amplitudes. Same format as panel D. *F*, Gradient-echo versus spin-echo. We re-plot results from panels D and E, directly comparing the two datasets acquired using gradient-echo against the two datasets acquired using spin-echo (conducted in the same subjects).

Next, we constructed depth profiles in order to understand how the magnitude of BOLD responses change with cortical depth. For these analyses, we restricted our quantification of BOLD responses to portions of cortex that have at least some substantive BOLD response for each given experimental condition. Specifically, for each condition, we selected vertices whose Standard beta exceeds 3% signal change at any cortical depth. We then averaged the BOLD response across these vertices at each depth (thus, a common set of vertices is used across depths). Consistent with earlier inspections of surface maps, we find that the magnitudes of both Standard betas and Late betas exhibit strong depth dependence (**Figure 8D**). For example, the BOLD signal change for Late betas is about 4 times greater at Depth 1 (outer) than at Depth 6 (inner). In contrast, the BOLD signal change for Early betas is fairly constant across depth. This effect is observed both in the group average (**Figure 8D, main plot**) as well as in each individual dataset (**Figure 8D, inset plots**). Thus, TDM successfully removes late components, which likely reflect unwanted macrovascular effects that bias BOLD responses towards outer cortical depths.

Finally, an important aspect of betas is their reliability, i.e. robustness or consistency across repeated measurements. We observed earlier that the eccentricity maps vary substantially in quality (see **Figure 6, fifth through seventh columns**), suggesting that betas can suffer from low reliability. To quantify these observations, we calculated the correlation of betas across split-halves of each dataset (test-retest reliability). We find that the Standard betas are the most reliable, and there is a drop in reliability for the Early and Late betas (**Figure 8E**). The decrease in reliability for the Early and Late betas is unfortunate but not surprising: Early and Late timecourses are typically highly correlated, and the accuracy of regression estimates in the case of correlated regressors is expected to be somewhat degraded.

### 3.7. TDM is compatible with other acquisition methods

To gain further insight into the nature of TDM and its generalizability, we repeated the eccentricity experiment using low-resolution fMRI (3T, 2.4-mm; Datasets D15–D16) as well as spin-echo fMRI (7T, 1.05-mm; Datasets D13–D14). The acquisition was performed in subjects who also participated in the high-resolution gradient-echo acquisition, enabling direct comparison of results.

We find that the TDM method successfully applies to both acquisition styles. In the low-resolution data, we see that the distribution of timecourse shapes tightens (**Supplementary Figure 3, fifth and seventh rows**), which likely reflects a combination of averaging diverse timecourses within individual voxels and reduction of thermal noise. Moreover, we find that TDM derives Early and Late timecourses that closely resemble those found in the high-resolution data (**Supplementary Figure 3, fourth column**). This implies that even at a resolution of 2.4 mm, there is sufficient diversity of timecourses to support data-driven discovery of latent timecourses. Examining the surface visualizations (**Figure 7**), we observe results for the low-resolution measurements that are consistent with the high-resolution measurements. However, the differences between the spatial patterns of the Early and Late betas are reduced, indicating that the differences between microvascular and macrovascular effects tend to wash out at low spatial resolutions.

In the spin-echo data, we find patterns of results that look remarkably similar to the gradient-echo data. Early and Late timecourses are identified (**Supplementary Figure 3, first and third rows**), and large betas are found for Late timecourses (**Figure 7**). Standard GLM analysis of spin-echo data yields betas that exhibit depth-dependent bias in BOLD signal change (**Figure 8F, upper right, black lines**), and this bias is largely eliminated after applying TDM (**Figure 8F, upper right, red lines**). Importantly, the spin-echo measurements suffer from a decrease in sensitivity compared to the gradient-echo measurements (**Figure 8F, bottom plots, black lines**), even though larger voxels were used for the spin-echo measurements. Overall, these results indicate that spin-echo measurements still contain substantial contributions from the macrovasculature and that TDM is able to identify and remove the macrovasculature-related effects.

## 4. Discussion

### 4.1. Nature of the TDM method

The core idea underlying TDM is that early and late timecourses can be derived from task-based fMRI data and reflect microvasculature-related and macrovascular-related responses, respectively. This idea is conceptually simple and draws its roots from critical observations made in prior studies (de Zwart et al., 2005; Lee et al., 1995; Siero et al., 2011). The value of the work presented here lies not so much in the discovery of a new phenomenon, but rather in the design and validation of new analysis methodology. Specifically, we have designed analysis procedures that are simple and robust and produce readily interpretable visualizations and results. Furthermore, we provide substantial empirical validation that the technique, when applied to fMRI measurements, produces sensible results: late components co-vary with dark EPI intensity (**Figures 6–7, Supplementary Figure 2**), depth-dependent response profiles are substantially flattened after removal of late components (**Figure 6, Figure 8D**), and artifacts in cortical maps are eliminated (**Figures 6–7**).

The TDM method has a few prerequisites:

- *Data acquisition*. The TDM method is likely compatible with a broad range of acquisition styles. As we have shown, TDM can be applied to data from standard spatial resolutions (2–3 mm; see **Figure 7, Supplementary Figure 3**) or data from high spatial resolutions (<1 mm; see **Figure 6**). Presumably, the spatial resolution needs only to be high enough to allow diverse sampling of vasculature in the brain. With respect to temporal resolution, we have shown that TDM can be successfully applied to data acquired at even fairly slow rates, such as the 2.2-s sampling rate used in Datasets D1–D14; this was likely aided by the fact that we jittered the acquired time points with respect to the experimental conditions (see **Figure 1B**).
- *Experimental design*. The TDM method requires a task-based experiment in which neural events occur at prescribed times, and is therefore inapplicable to resting-state paradigms. We suspect that TDM will be most effective for event-related paradigms where experimental events are somewhat short (e.g. 4 s or less). Block designs involving prolonged events (e.g. 16–30 s) or designs involving continuously changing experimental parameters (e.g. sinusoidal variation of a stimulus property) are likely to generate microvasculature-related and macrovascular-related timecourses that are more similar and therefore harder to disambiguate.
- *Amount of data*. Because TDM is a data-driven technique, it is necessary to acquire sufficient data to support the method. For example, there must be sufficient data to estimate response timecourses from the voxels in a dataset. If a given dataset is overly noisy or if not enough data are collected, timecourse estimates may be noisy and nearly isotropic in their distributions in the 3-dimensional PCA space (e.g. see Dataset D6 in **Supplementary Figure 2**), making it difficult to extract the underlying manifold structure of the data.

Given that these prerequisites are fairly minimal (i.e., event-related designs with reasonable fMRI acquisition parameters and reasonable amounts of data), we suspect that the TDM method may be widely applicable to different kinds of fMRI data.

Algorithmically, TDM uses a manifold-fitting method to characterize latent structure in timecourse variations. There are other methods that can characterize latent structure; two widely used methods are principal components analysis (PCA) and independent components analysis (ICA). Could these methods have been used instead? Due to the orthogonality constraint in PCA, it is necessarily the case that the PC timecourses returned by PCA are orthogonal (i.e. the dot product between each pair of timecourses equals zero). Although TDM does make use of PCA to determine the 3-dimensional space within which to perform further analyses, the PCA timecourses themselves do not constitute good candidates for latent timecourses. This is simply because there is no reason to expect hemodynamic timecourses in the brain to be orthogonal. Indeed, the bulk of empirically measured timecourses tend to reside in a small portion of the 3-dimensional space, and the early and late timecourses returned by TDM are nearby in this space and highly correlated (see **Figure 4** and **Supplementary Figure 2**).

In contrast to PCA, ICA does not impose the constraint of orthogonality. Instead, ICA optimizes timecourses with respect to statistical independence, often through some measure of non-Gaussianity (e.g. kurtosis). We have demonstrated that it is possible to construct an ICA-based procedure that can potentially derive early and late timecourses. Though the derived timecourses from the ICA-based procedure are sometimes similar to those produced by TDM (see x’s in **Supplementary Figure 2**), there are clear advantages of the TDM method. First, the data visualization and explicit modeling performed by TDM allow the user to evaluate and confirm the data features that give rise to the derived timecourses. ICA, without further analysis, remains a ‘black box’ and it is difficult to understand the specific features of the data that give rise to its results. Second, there is no *a priori* reason to think that loadings on early and late hemodynamic timecourses must necessarily conform to statistical independence. Thus, relying on a procedure that is predicated on independence seems risky. Third, ICA alone does not identify the early and late timecourses; rather, we found it necessary to couple the results of ICA with several post-hoc procedures that are somewhat heuristic in nature and thus unsatisfying (see Section 2.8.4). On the whole, we suggest that TDM is more explicit, more direct, and more interpretable than ICA. Indeed, historically, the first method that we developed was the ICA-based procedure, and the shortcomings described above are what prompted us to develop the TDM method.

GLM-based analyses of fMRI time-series data sometimes allow flexible modeling of timecourse shape through the inclusion of a canonical hemodynamic response timecourse and its temporal derivative (Friston et al., 1998a) or some other basis function decomposition such as PCA (Woolrich et al., 2004). While TDM shares the common feature of providing a means to capture timecourse variation, the key difference with respect to these alternative approaches lies in the specific timecourses that are chosen by TDM. The Early and Late timecourses found by TDM are often quite correlated (unlike a timecourse and its derivative or those returned by PCA). Moreover, the Early and Late timecourses have specific biological meanings, and so the beta loadings found for these timecourses have specific value. It is possible that alternative timecourse models can yield fits to a set of data that are as good as the fit achieved by TDM, but the beta loadings associated with these models cannot be interpreted in terms of microvasculature- and macrovasculature-related components.

### 4.2. Validation of the TDM method

In this study, we demonstrated that TDM delivers robust and meaningful results in each of the 16 fMRI datasets collected (11 unique subjects). The main lines of validation include sensible timecourses (the shapes of the Early and Late timecourses are plausible and consistent with simple inspections of response timecourses; see **Figure 2, Supplementary Figure 1**), co-variation of loadings on the Late timecourse with dark EPI intensities (see **Figure 6**) and with kurtotic BOLD amplitudes (see **Figure 8B**), flattening of depth-dependent response profiles (see **Figure 8D**), and elimination of artifacts in cortical maps for which we have ground-truth expectations (see **Figures 6–7**). Furthermore, we make available data and analysis code to ensure that the TDM method is transparent and reproducible (Poline and Poldrack, 2012).

While we believe a solid case has been made for TDM, additional validation would nonetheless be useful. Further work could be directed at assessing and optimizing the technique with respect to experimental design characteristics such as the duration of experimental conditions, the spatial and temporal resolution of the acquisition, and the amount of data acquired. In addition, it would be worthwhile to test the technique on other types of experiments (other sensory, cognitive, and/or motor experiments) and in other brain areas. In particular, it would be interesting to assess how well TDM can resolve fine-scale variation in neural representations, such as ocular dominance columns (Cheng et al., 2001). Since TDM makes no restrictions on the spatial loadings of the Early and Late timecourses, the technique should in principle be applicable not only to large-scale neural representations like eccentricity but also fine-scale representations like ocular dominance. Finally, it would be quite valuable to validate results obtained using TDM against direct neural activity measurements, such as electrophysiological recordings with laminar resolution (Maier et al., 2010; Self et al., 2019).

### 4.3. Strengths and limitations of the TDM method

All methods have strengths and limitations. To make an informed decision of whether to pursue a given method, it is important to understand its strengths and limitations.

Strengths of the TDM method include the following:

- *Flexibility*. As discussed in Section 4.1, TDM has fairly minimal prerequisites. It can be applied to different types of acquisitions: gradient-echo, spin-echo, high-resolution, low-resolution, etc. Moreover, as a data-driven method, TDM makes no assumptions about shapes of hemodynamic timecourses that might be found in a dataset and naturally adapts to different brain areas, subjects, and/or datasets. Furthermore, since the response decomposition is performed independently for each voxel, the technique makes no assumptions regarding the spatial distribution of BOLD responses in a given experiment. In other words, the loadings on the Early and Late timecourses can vary from voxel to voxel, and the technique can, in theory, capture these variations. Finally, TDM applies to single-condition activity maps, and therefore avoids the assumptions required in differential paradigms where unwanted non-specific effects are assumed to be removed through subtraction.
- *Transparency*. An integral aspect of TDM is direct visualization of distributions of timecourses. Thus, it is easy for the user to understand the nature of the data and whether the derived timecourses are meaningful. This stands in contrast to ‘black box’ methods that might produce unusual results without explanation.
- *Simplicity and robustness*. TDM is simple in its design and does not, as far as we have seen, require fine-tuning of parameters to be effective. Our results establish more than just a proof of principle: we show that the method without any modification performs robustly across different datasets, subjects, and experiments.
- *Analysis not acquisition*. Since TDM is an analysis method, it can be retrospectively applied to datasets that have already been acquired. Moreover, since TDM does not place major constraints on acquisition, the user is not burdened with making difficult decisions regarding optimal acquisition parameters (e.g., choosing between a standard acquisition scheme that is guaranteed to produce reasonably strong signals versus a specialized acquisition scheme that might suffer from low sensitivity).

Limitations of the TDM method include the following:

- *Sensitivity loss*. TDM involves decomposing fMRI responses using two timecourses that are often highly correlated. Thus, from a regression perspective, one expects to incur a penalty in terms of high variance in beta estimates. One can view the use of TDM as approximately doubling the number of experimental conditions while keeping the number of data points constant. As such, the ensuing model will be more difficult to reliably estimate compared to a basic GLM model with a single set of experimental conditions. Thus, if sensitivity (i.e. reliability of beta estimates) is the sole priority and specificity (i.e. accurate estimates of local neural activity) is not critical, the TDM method is not recommended. If, on the other hand, specificity is of utmost importance, TDM is likely to be a valuable method. In short, TDM does not deliver more robust fMRI maps (e.g. large blobs of statistically significant activations), but aims to deliver more spatially accurate and neurally meaningful maps.
- *Intrinsic physiological limitations*. The TDM method attempts to disambiguate BOLD contributions from the microvasculature and macrovasculature based on their respective associated timecourses. The more similar these timecourses, the more difficult it will be to estimate the distinct contributions of the timecourses. Thus, the intrinsic physiology of the subject places limits on the overall effectiveness of the TDM method. For example, in our data (see **Supplementary Figure 2**), we find that Subject S5 exhibits Early and Late timecourses that are widely separated in the 3-dimensional PCA space and are highly distinct, whereas Subject S4 exhibits Early and Late timecourses that are close together in the 3-dimensional PCA space and are fairly similar. This may be the reason why some datasets (such as Subject S4’s Dataset D4) experience a larger reduction in reliability when using TDM compared to other datasets (see **Figure 8E**). One possible approach to achieve optimal results is to screen subjects according to the temporal separability of their microvasculature- and macrovasculature-related timecourses.
- *Complexity of the vasculature*. TDM proposes a simple two-component model to decompose BOLD timecourses. The vasculature is certainly more complex than this simple characterization (Uludağ and Blinder, 2018). Thus, it may be fruitful to develop a more nuanced characterization of vasculature types and their dynamics.

### 4.4. Comparison to other methods

How does TDM compare to other approaches for removing or avoiding venous effects in fMRI? Some researchers have proposed simple heuristic selection methods. For example, sampling BOLD responses at only deep cortical depths (Polimeni et al., 2010) can help avoid the influence of large draining veins near the pial surface. However, this comes at the cost of not being able to infer response properties in superficial cortical depths; moreover, it is still possible for veins to penetrate deep into cortex (see **Figure 6**). Another example is masking out voxels with very high percent BOLD signal change (e.g. Shmuel et al., 2007), low EPI intensity (e.g. Olman et al., 2007), and/or low temporal signal-to-noise ratio (e.g. Fracasso et al., 2018). While these are reasonable heuristics for removing voxels that are most egregiously affected by large veins, it is not clear what principle can be used to set the threshold to be used. Moreover, similar to the approach of sampling only deep cortical depths, this approach fails to recover usable signals from the removed voxels. Finally, one suggestion found in older work (Goodyear and Menon, 2001; Shmuel et al., 2007) and more recent work (Blazejewska, Nasr, Polimeni, ISMRM 2018 abstract) is to sample early time points in the BOLD response. This is certainly consistent with the spirit of TDM and may produce a response snapshot that is more weighted towards the microvasculature. However, our results indicate that Early and Late timecourses are highly overlapping (see **Figure 5**). If the chosen time point is not sufficiently early, this incurs the risk of the late component “bleeding” into the analysis results. Moreover, choosing only one or a few time points does not make efficient use of all of the available data. Compared to these various heuristic selection methods, we believe that TDM has substantial appeal: *TDM can recover signals at all depths and even in voxels that have substantial venous influence; it makes efficient use of all of the fMRI data collected; and, as a data-driven technique, it naturally identifies appropriate timecourse parameters for each dataset*.

Recently, a method has been proposed that first constructs a forward model characterizing the mixing of hemodynamic signals from different cortical depths due to blood drainage towards the pial surface and then uses this model to invert observed BOLD depth profiles (Heinzle et al., 2016; Markuerkiaga et al., 2016; Marquardt et al., 2018). This method bears a parallel to TDM in the sense that it is a spatial deconvolution approach whereas TDM is a temporal decomposition approach. However, the accuracy of the spatial deconvolution approach may be dependent on the correctness of the model parameters, which might vary across brain regions and/or subjects. In addition, the approach deals only with vascular effects that vary across depths. In contrast, TDM is a data-driven technique that adapts to each given dataset and compensates for vascular effects present at every voxel.

Besides analysis methods, one can consider using acquisition methods to avoid venous effects. Switching from conventional gradient-echo pulse sequences to spin-echo pulse sequences (or related techniques such as GRASE (De Martino et al., 2013; Moerel et al., 2018; Olman et al., 2012)) has traditionally been considered the standard approach for mitigating venous effects in fMRI (Yacoub et al., 2008). While the refocusing of T2* effects by the 180° RF pulse in spin-echo eliminates sensitivity to extravascular effects around large veins, it is important to note this holds only for a specific point in time (typically the center of the readout window). The remainder of the image acquisition incurs T2* effects (Goense and Logothetis, 2006). Furthermore, spin-echo does not eliminate intravascular effects in large vessels (Budde et al., 2014; Duong et al., 2003). Thus, spin-echo does not provide full elimination of venous effects. In addition, spin-echo acquisitions suffer from increased energy deposition, limits on spatial coverage, lower temporal resolution, and loss of signal-to-noise ratio.

In this study, we have provided a direct comparison of TDM and spin-echo. In order to maintain sensitivity, we acquired spin-echo data at a lower resolution (1.05-mm vs. 0.8-mm) but maintained the same TR and the same overall experiment duration as the gradient-echo data. Our results show that the spin-echo data analyzed using a standard GLM (single canonical hemodynamic timecourse) is more robust than gradient-echo data decomposed using TDM (see **Figure 8F**). One reason for the increased robustness of the spin-echo data is its lower spatial resolution, providing the data with some advantage over the gradient-echo data. However, even if the two types of data were matched in resolution, spin-echo should not be viewed as a complete solution since it does not fully suppress venous effects. Indeed, we demonstrate that TDM can be applied to the spin-echo data in order to remove venous influences present in those data (see **Figure 7**). When comparing gradient-echo data and spin-echo data that have both been decomposed using TDM, gradient-echo has greater robustness (see **Figure 8F**). Thus, we suggest that if removal of venous effects is top priority and one plans to use TDM, there is little benefit to spin-echo acquisition over conventional gradient-echo acquisition.

A promising alternative to spin-echo is vascular space occupancy (VASO), a pulse sequence that is not sensitive to the BOLD effect but rather to changes in cerebral blood volume (Lu et al., 2003). This approach avoids the impurities that linger in spin-echo acquisitions, and has been shown to generate highly specific measures of task-driven hemodynamic responses (Huber et al., 2017). A promising direction for future work is to perform direct comparison of an gradient-echo acquisition optimized for use with TDM against an optimized VASO acquisition. Through rigorous evaluations, we hope that the field will eventually reach consensus with respect to the most effective approach for achieving accurate fine-scale measurements of brain activity.

## 5. Author Contributions

K.K. designed the experiment. R.Z. collected the data. K.K. and K.J. developed techniques and analyzed the data. K.K. wrote the paper. All authors discussed and edited the manuscript.

## 6. Acknowledgements

We thank E. Margalit and N. Petridou for helpful discussions. This work was supported by NIH Grants P41 EB015894, P41 EB027061, P30 NS076408, S10 RR026783, S10 OD017974-01, U01 EB025144, and the W. M. Keck Foundation.

## Supplementary Material

**Supplementary Figure 1.**
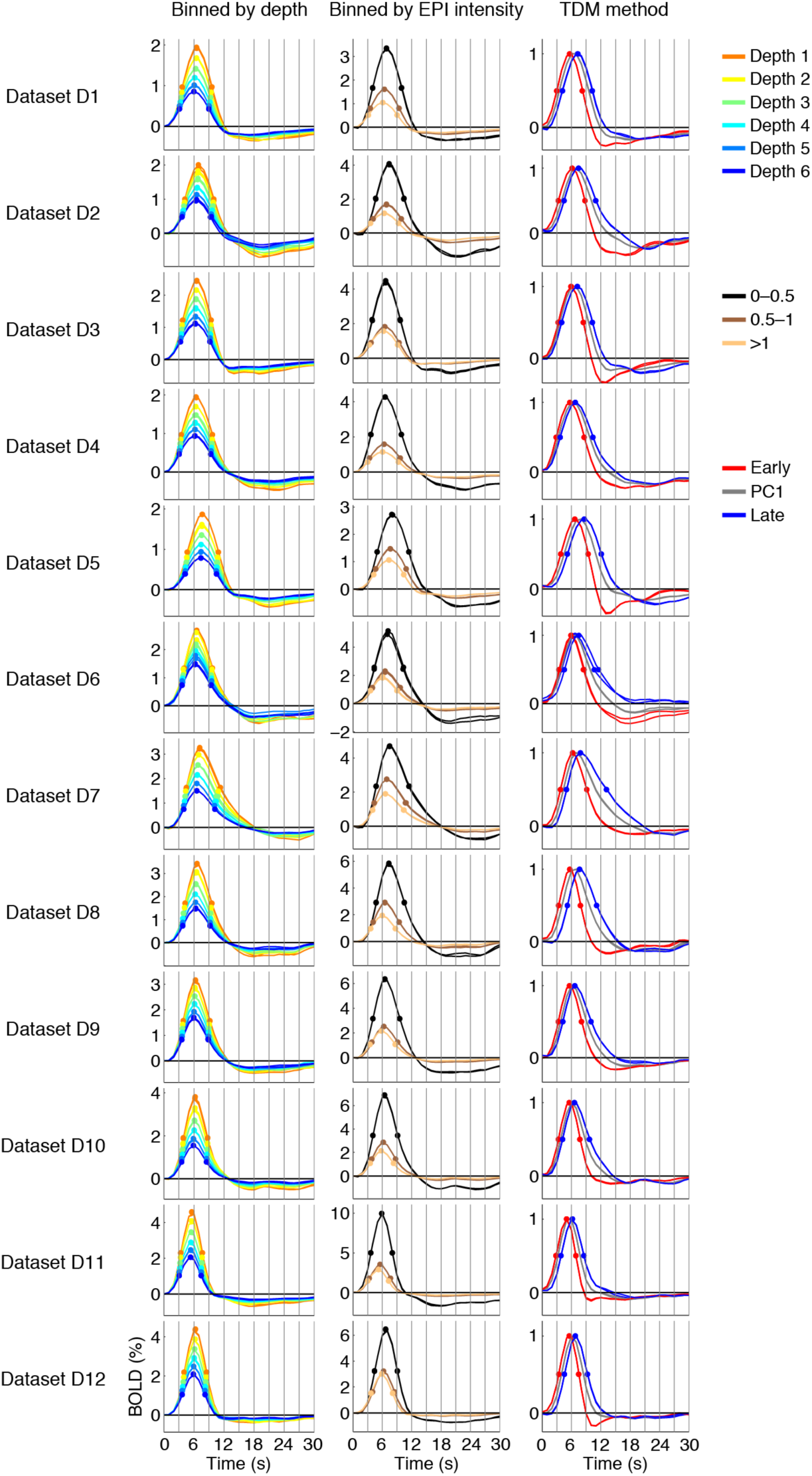
Comprehensive summary of BOLD timecourses. Same format as Figure 2. Qualitative patterns of results are highly consistent across datasets (e.g. time-to-peak is delayed at superficial depths). However, there is substantial quantitative variation across datasets (e.g. time-to-peak is short in Dataset D11 but long in Dataset D5). This underscores the importance of tailoring timecourse derivation to individual subjects or scan sessions.

**Supplementary Figure 2.**
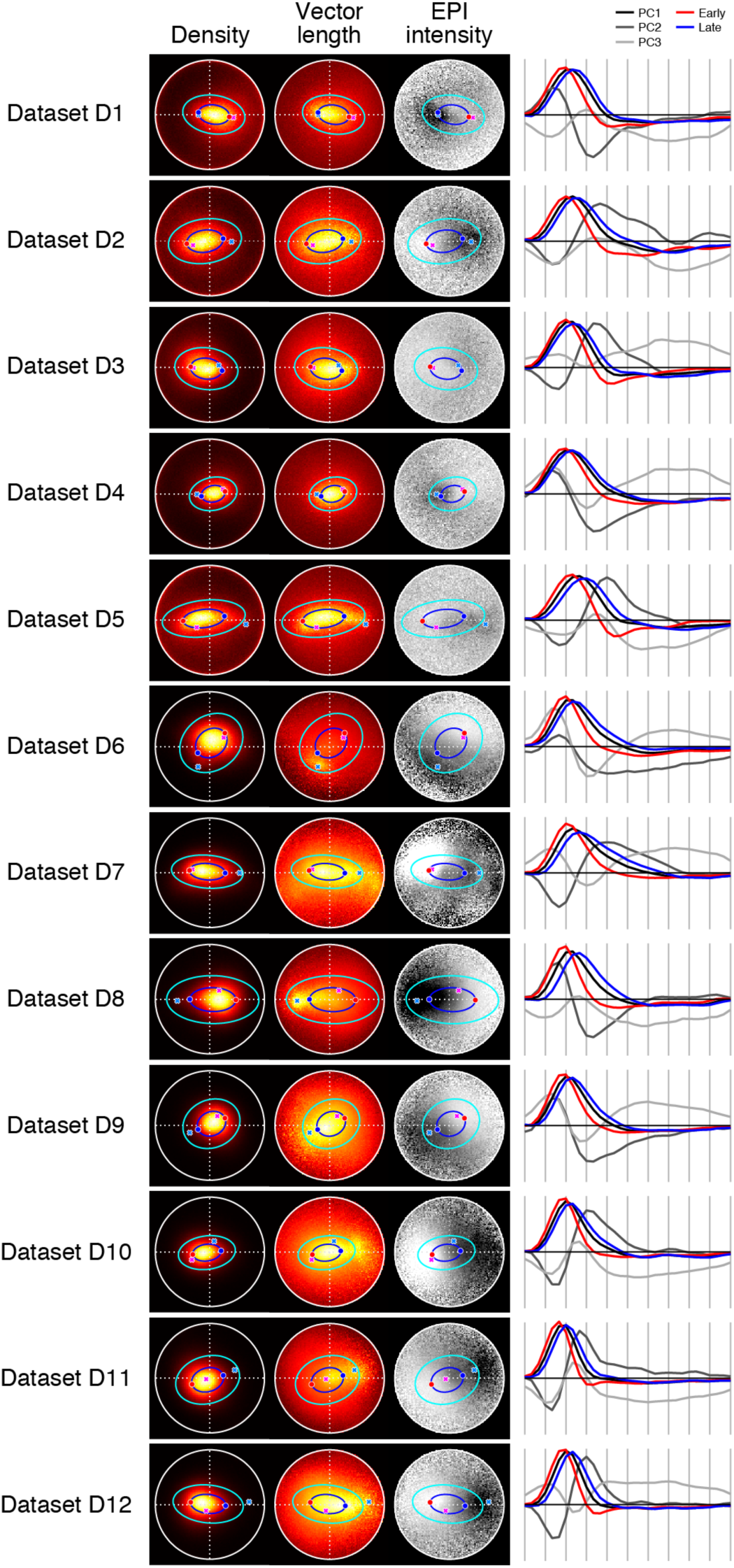
TDM results for the high-resolution gradient-echo datasets (D1–D12). Same format as Figure 4, except for the following addition: magenta and cyan crosses indicate the early and late timecourses derived from the ICA-based procedure (see Section 2.8.4). TDM consistently identifies reasonable early and late timecourses in each dataset. The ICA-based procedure yields similar timecourses in some datasets (e.g. D4), but diverges substantially in others (e.g. D8). It appears that timecourses with very large BOLD responses (see D8) is a major factor that influences the timecourses returned by ICA.

**Supplementary Figure 3.**
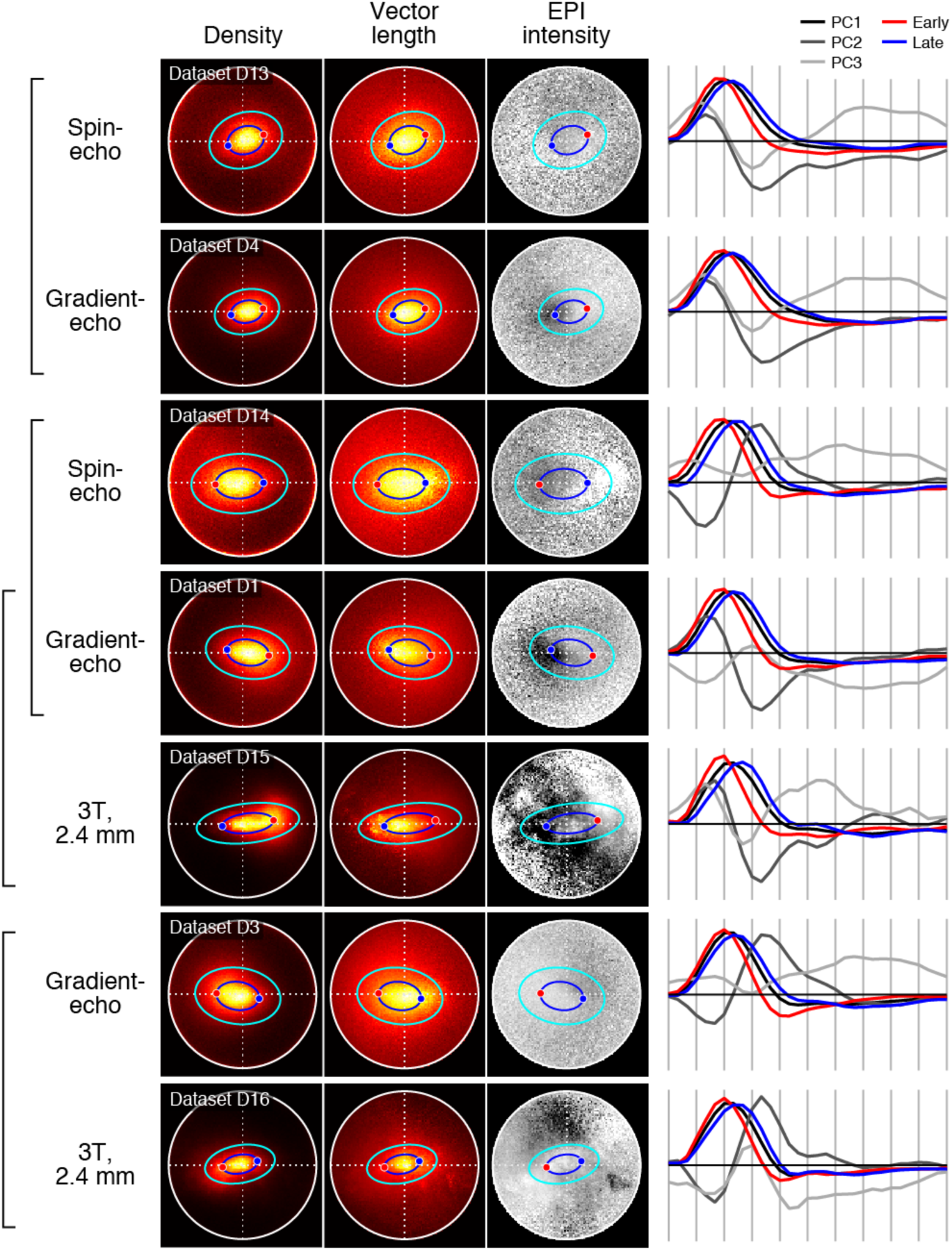
TDM results for the alternative acquisition protocols (D13–D16). Same format as Figure 4. To facilitate comparison, we place results obtained using the spin-echo and low-resolution protocols next to results obtained using the high-resolution gradient-echo protocol.

